# Preclinical models of human multiple myeloma subgroups

**DOI:** 10.1101/2021.08.28.458010

**Authors:** Wiebke Winkler, Carlota Farré Díaz, Eric Blanc, Hanna Napieczynska, Patrick Langner, Marvin Werner, Barbara Walter, Brigitte Wollert-Wulf, Tomoharu Yasuda, Arnd Heuser, Dieter Beule, Stephan Mathas, Ioannis Anagnostopoulos, Andreas Rosenwald, Klaus Rajewsky, Martin Janz

## Abstract

Multiple myeloma (MM), a tumor of germinal center (GC)-experienced plasma cells, comprises distinct genetic subgroups, such as the t(11;14)/CCND1 and the t(4;14)/MMSET subtype. We have generated subgroup-specific MM models by the GC B cell-specific co-activation of Ccnd1 or MMSET with a constitutively active Ikk2 mutant, mimicking the secondary NFκB activation frequently seen in human MM. Ccnd1/Ikk2ca and MMSET/Ikk2ca mice developed a pronounced, clonally restricted plasma cell outgrowth with age, accompanied by serum M spikes, bone marrow insufficiency and bone lesions. The transgenic plasma cells could be propagated *in vivo* and showed transcriptional profiles resembling their human counterparts. Thus, we show that Ccnd1 and MMSET cooperate with NFκB in MM pathogenesis, considering for the first time the genetic heterogeneity of MM for the generation of preclinical models.

## Main Text

Multiple myeloma (MM) is a tumor of antibody-producing plasma cells that expand within the bone marrow (BM). The disease shows a stepwise progression from a pre-malignant stage (monoclonal gammopathy of undetermined significance, MGUS) to overt MM, commonly defined by the occurrence of CRAB symptoms (i.e. hyper<u>c</u>alcemia, <u>r</u>enal failure, <u>a</u>nemia, <u>b</u>one lesions) ((*1*), (*2*)). Recurrent early, MGUS-initiating genetic events include chromosomal translocations into the immunoglobulin heavy chain (IGH) locus ((*3*), (*4*), (*5*)), which typically arise during the germinal center (GC) reaction ((*6*), (*7*)) and frequently affect CCND1, t(11;14), or the histone methyltransferase MMSET, t(4;14), in 15-20 % and 15 % of MM cases, respectively ((*8*), (*9*)). The MGUS to MM transition requires additional genetic aberrations, such as c-MYC translocations, aberrant NFκB or MAPK signaling and p53 lesions ((*10*)). This complex genetic architecture characterizes MM as a heterogeneous disease, in line with gene expression studies that separate MM patients into 7-10 distinct subgroups ((*11*), (*12*), (*13*)). Of these, four are associated with a particular IGH translocation, e.g. t(4;14) with the MMSET group, implying an important role of the early genetic aberrations for tumor biology and sub-classification. Given that the distinct MM subgroups differ in terms of gene expression, tumor biology and clinical outcome, new therapeutic strategies have to take into account the heterogeneity of the disease. One approach to delineate subgroup-specific disease mechanisms is the creation of subgroup-specific animal models. However, the existing transgenic MM models are not suitable for this analysis as they focus on progression-associated events like c-MYC ((*14*)), plasma cell-related genes such as XBP1 ((*15*)), signaling molecules like gp-130 ((*16*)) and/or do not comply with the GC origin of the initiating events ((*17*), (*18*)).

With the aim to establish subgroup-specific MM models for the two most frequent primary translocation events, t(11;14) and t(4;14), we generated conditional gain-of-function alleles for mouse Ccnd1 and MMSET in the Rosa26 locus (Fig. 1A-B, S1). To mimic the secondary event necessary for MGUS-MM transition, we combined these alleles with the R26 Ikk2ca^stopF^ allele ((*19*)) that expresses a constitutively active Ikk2 mutant, mimicking the NFκB activation commonly seen in human MM, either due to genetic aberrations ((*20*), (*21*)) or BM microenvironment-mediated signaling ((*22*)) (Fig. 1A-B). To restrict transgene activation *in vivo* to GC B and plasma cells, R26 Ccnd1^stopF^ (hereafter termed Ccnd1) and R26 MMSET^stopF^ (hereafter: MMSET) and R26 Ikk2ca^stopF^ (hereafter: Ikk2ca) mice were crossed to the C*γ*1-cre strain ((*23*), Fig. 1B). Notably, all single (Ccnd1, MMSET or Ikk2ca) and double (Ccnd1/Ikk2ca or MMSET/Ikk2ca) mutant mice were able to mount normal GC and antibody responses upon immunization with the T cell-dependent antigen NP-CGG (4-hydroxy-3-nitrophenylacetyl-chicken-gamma-globulin) (Fig. S2). To analyze whether the GC B cell-specific activation of Ccnd1 or MMSET alone or in combination with Ikk2ca results in MM development, mice were immunized once and then aged to monitor tumor development (Fig. 1B). Within our t(4;14) cohort, MMSET/Ikk2ca mice reached disease-defining end-points between 72-97 weeks of age, presenting with splenomegaly and porous long bones (Fig. S3A, S4A). HE staining and immunofluorescence analyses of femur sections revealed a pronounced expansion of large, immunoglobulin light chain positive plasma cells with abundant cytoplasm and eccentric nuclei in the vast majority of MMSET/Ikk2ca samples, e.g. 60 % infiltration in 10 of 15 mice, compared to normal femur morphology in most age-matched control and single mutant mice (Fig. 1C, Table S1). Similarly, plasma cells dominated in MMSET/Ikk2ca spleens (Fig. S3G, Table S1). Correspondingly, MMSET/Ikk2ca mice showed a significant increase of spleen and BM CD138+TACI+ plasma cells by flow cytometry, most of which expressed both reporters, i.e. BFP labeling MMSET and GFP marking Ikk2ca expression, suggestive of a synergistic effect of the two transgenes (Fig. S3C-F). However, the percentage of plasma cells determined by flow cytometry was lower than that observed in histology (Fig. S3D), most likely reflecting rapid cell death *ex vivo*, consistent with a strong overall reduction of recovered cells (Fig. S3B). The observed plasma cell expansion was accompanied by elevated immunoglobulin serum titers (Fig. S3H) and an enrichment of antibody-secreting BM cells (Fig. S3I). Importantly, a high proportion of MMSET/Ikk2ca mice developed pathologies associated with MM. The most prominent features were increased serum calcium levels (> 9.6 mg/dL; 11/15 mice), anemia (hemoglobin < 10.2 g/dL; 6/15 mice; red blood cells < 6.8 M/µL; 11/15 mice), thrombocytopenia (platelets < 662 K/µL; 14/15 mice), and hypoalbuminemia (< 23 g/L; 10/15 mice) (Fig. 1D, S4C-D). Moreover, a substantial fraction of MMSET/Ikk2ca mice showed lesions within the tibia (8/16 mice) and extended areas of qualitatively lower bone density in the skull, reminiscent of osteopenia (6/16 mice) (Fig. 1E-F, S4A-B, S4D), indicating bone involvement.

**Figure 1:**
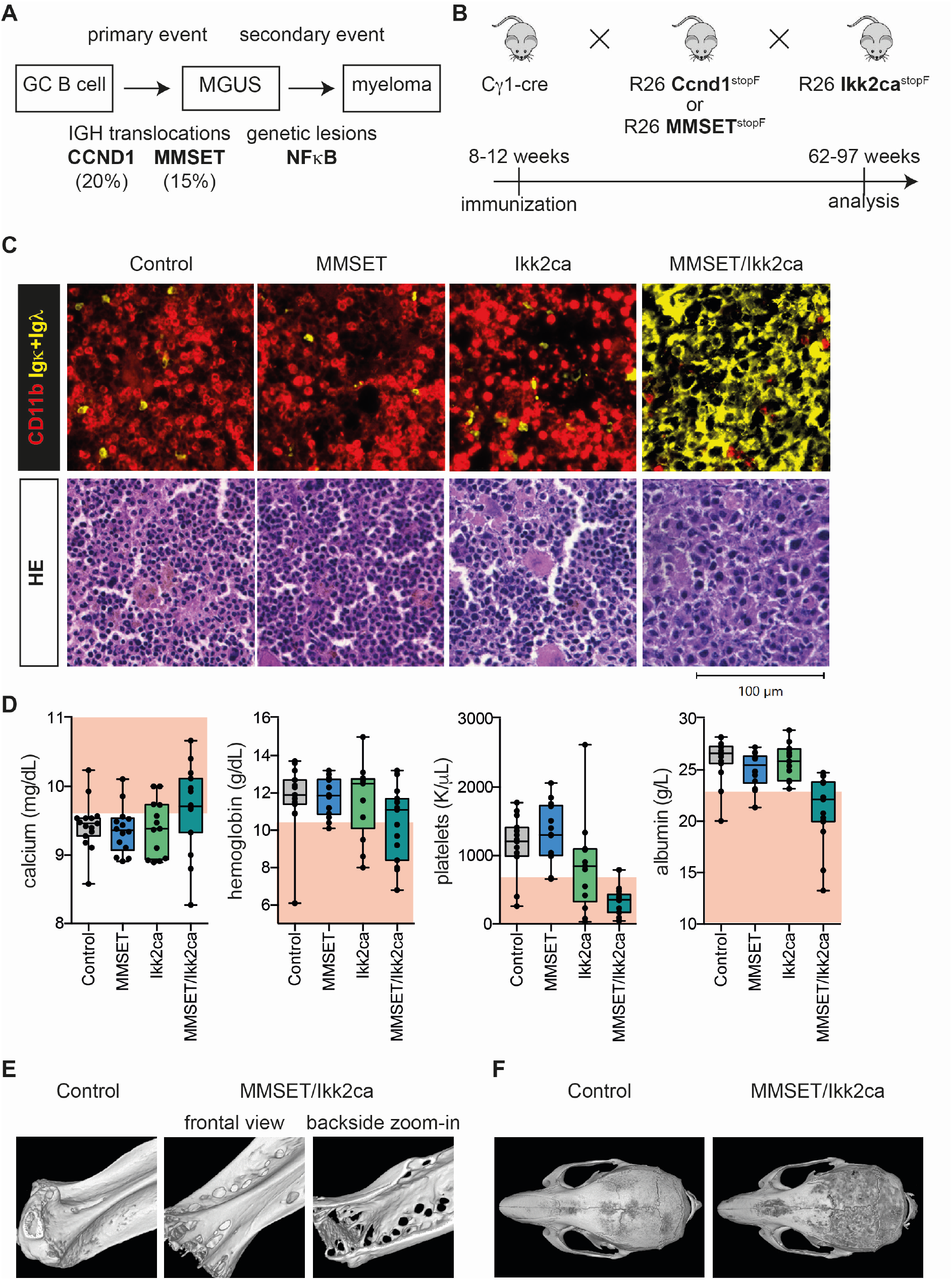
MMSET/Ikk2ca mice develop a plasma cell expansion with age. (A) Simplified scheme of multistep human MGUS-MM development, illustrating the sequential acquisition of initiating (t(11;14)/CCND1 or t(4;14)/MMSET) and secondary (NFκB activation) genetic lesions. (B) Outline for the generation of MM subgroup-specific mouse cohorts by the GC B cell-specific activation of MM-associated factors using the Cγ1-cre strain. Control, single (Ccnd1, MMSET or Ikk2ca) and double mutant (Ccnd1/Ikk2ca or MMSET/Ikk2ca) mice were immunized with NP-CGG at 8-12 weeks of age, monitored for tumor development and sacrificed upon development of disease-defining endpoints (between 62-97 weeks of age). (C) Upper panel: representative immunofluorescence images of femur sections stained with α-CD11b (red, myeloid cells) and α-Igκ/α-Igλ (yellow, plasma cells). Lower panel: representative HE stained femur sections. (D) Graphs depicting serum calcium, hemoglobin, platelets and serum albumin values of MMSET cohort mice. The pathologic range of either parameter (calcium >9.6mg/dL; hemoglobin <10.2g/dL; platelets <662K/µL; albumin* <23g/L) is marked in red (reference: Jackson Laboratory; 78 weeks old C57BL/6J; *the lower limit for albumin had to be adjusted for our cohort to discern healthy and diseased mice for animals 78 weeks). (E-F) Representative 3D images reconstructed from µCT scans of tibiae (E) and skulls (F). Tibiae are shown in frontal view (E; left) and as zoomed-in image (E; right) to demonstrate that lesions destroyed the cortex.

A hallmark of MM is the accumulation of a monoclonal (M) immunoglobulin in the serum. In line with this, we detected prominent serum M spikes in MMSET/Ikk2ca mice compared to absent or only faint bands in control and single mutant mice (Fig. 2A). Subsequent immunofixation revealed the occurrence of one to three clones, mostly of the IgG or IgM isotype, suggestive of a clonally restricted plasma cell outgrowth (Fig. 2B, S7A). This finding was further supported by a limited Ighv gene repertoire in sorted double mutant CD138+TACI+ BM plasma cells. In the majority of MMSET/Ikk2ca mice, the respective dominant clone represented ≥ 50 % of all Ighv reads identified by RNA-sequencing (Fig. 2C, S5). Given our observation that a substantial fraction of MMSET/Ikk2ca plasma cells dies during *ex vivo* purification, the Ighv gene analysis of sorted cells probably underestimates the actual clonal restriction. The sorted population, however, is still representative of the *in vivo* situation as we reliably identified shared clones between sorted cells and femur or spleen sections by RNA-Seq and VDJ-PCR ((*24*)) (Fig. S5, S6, Table S2), as well as matching immunoglobulin isotype expression patterns of sorted cells and the respective serum M proteins (Fig. S7). To test whether the expanded plasma cells can be propagated *in vivo*, total spleen cells of a MMSET/Ikk2ca donor mouse were transplanted into Rag2-Il2rg-mice (Fig. 2D). Within 34 weeks, recipients presented with a prominent plasma cell outgrowth (Fig. 2E, S8) and serum M spikes (Fig. 2F). Strikingly, whereas the donor mouse demonstrated three different serum M proteins in immunofixation (Fig. 2B, #3939), 3 of 4 recipient mice showed only one major clone (Fig. 2G), implying increased clonal selection and accelerated expansion of transferred MMSET/Ikk2ca plasma cells.

**Figure 2:**
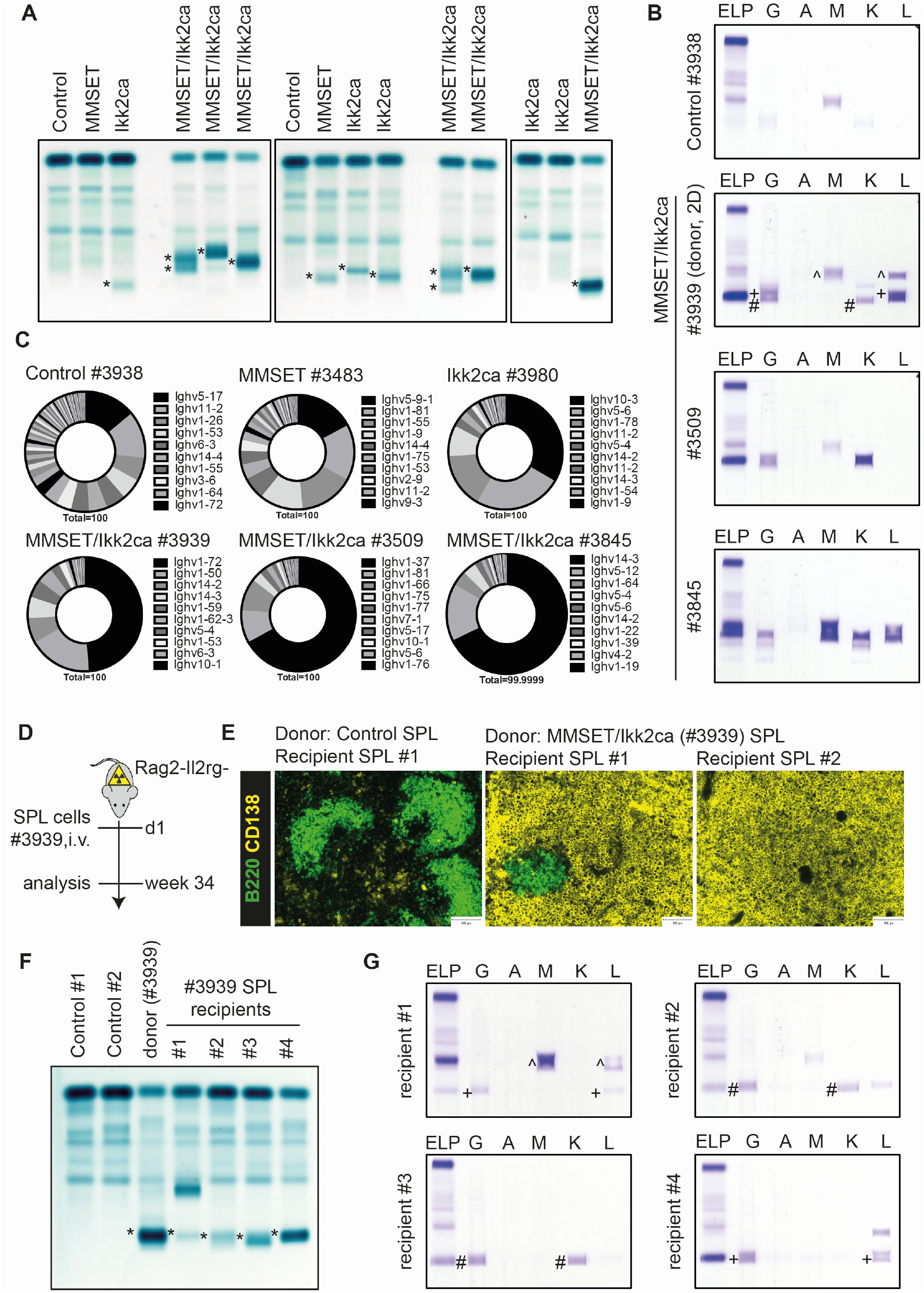
MMSET/Ikk2ca plasma cells are clonally restricted and can be propagated *in vivo*. (A) Representative serum protein electrophoresis (SPEP) of MMSET cohort mice; M spikes are marked with an asterisk. (B) SPEP coupled with immunofixation to determine the immunoglobulin isotype of the detected M proteins. (C) Ighv gene usage in sorted BM plasma cells of MMSET cohort mice. Pie charts show the fraction of individual Ighv genes among all annotated Ighv gene reads as determined by RNA-sequencing. The 10 most dominantly expressed Ighv genes are listed (ordered clockwise). (D) Scheme illustrating the total spleen cell transfer of either control or MMSET/Ikk2ca (#3939) mice into irradiated Rag2-Il2rg-recipients, followed by analysis after 34 weeks. (E) Representative immunofluorescence images of recipient spleens stained with α-B220 (green, B cells) and α-CD138 (yellow, plasma cells. (F) SPEP of two recipients transplanted with control splenocytes (Control #1+2), the MMSET/Ikk2ca donor (#3939) and its corresponding recipients; M spikes are marked with an asterisk. (G) SPEP coupled to immunofixation of the four MMSET/IKk2ca recipients to determine the immunoglobulin isotypes of the M proteins (corresponding heavy and light chains are marked by the same symbol; G - IgG, A - IgA, M - IgM, K - kappa, L - lambda).

Similar to MMSET/Ikk2ca mice, Ccnd1/Ikk2ca animals also developed a pronounced plasma cell phenotype between 62-91 weeks of age (Fig. 3, S9). The plasma cell outgrowth was confirmed by histology of femur and spleen sections demonstrating ≥ 50 % BM plasma cells in 6 of 7 double mutant mice (Fig. 3A, S9G, Table S3), increased immunoglobulin serum titers (Fig. S9H) and enriched antibody-secreting BM cells (Fig. S9I). As for MMSET/Ikk2ca mice, flow cytometry confirmed a significant expansion of transgene-expressing (BFP+GFP+) CD138+TACI+ cells, but greatly underestimated their frequency compared to histology (Fig. S9C-F). With regard to clinical parameters, Ccnd1/Ikk2ca mice also presented with hypercalcemia (4/6 mice), anemia (2/5 mice), thrombocytopenia (5/5 mice) and hypoalbuminemia (2/4 mice) (Fig. 3B, S10B-C). Likewise, bone disease was observed in two of six Ccnd1/Ikk2ca mice exhibiting tibial lesions, one of which also showed skull osteopenia (Fig. 3C, S10A, S10C). In addition, Ccnd1/Ikk2ca mice developed multiple M spikes of various Ig isotypes (Fig. 3D, S13), indicating an oligoclonal plasma cell outgrowth, which was further confirmed by a restricted Ighv gene repertoire by RNA-Seq (Fig. 3E, S11) and the detection of dominant clones by VDJ-PCR ((*24*)) (Fig. S12, Table S4). Ccnd1/Ikk2ca plasma cells could also be propagated upon total spleen cell transfer into immunodeficient hosts, with a massive plasma cell expansion within 31 weeks (Fig. 3F, S14A-B). Importantly, also in this setting the transferred plasma cells demonstrated increased clonal selection as only one dominant M spike could be detected in all recipients, whereas the donor mouse had presented with three (Fig. 3G, S14C).

**Figure 3:**
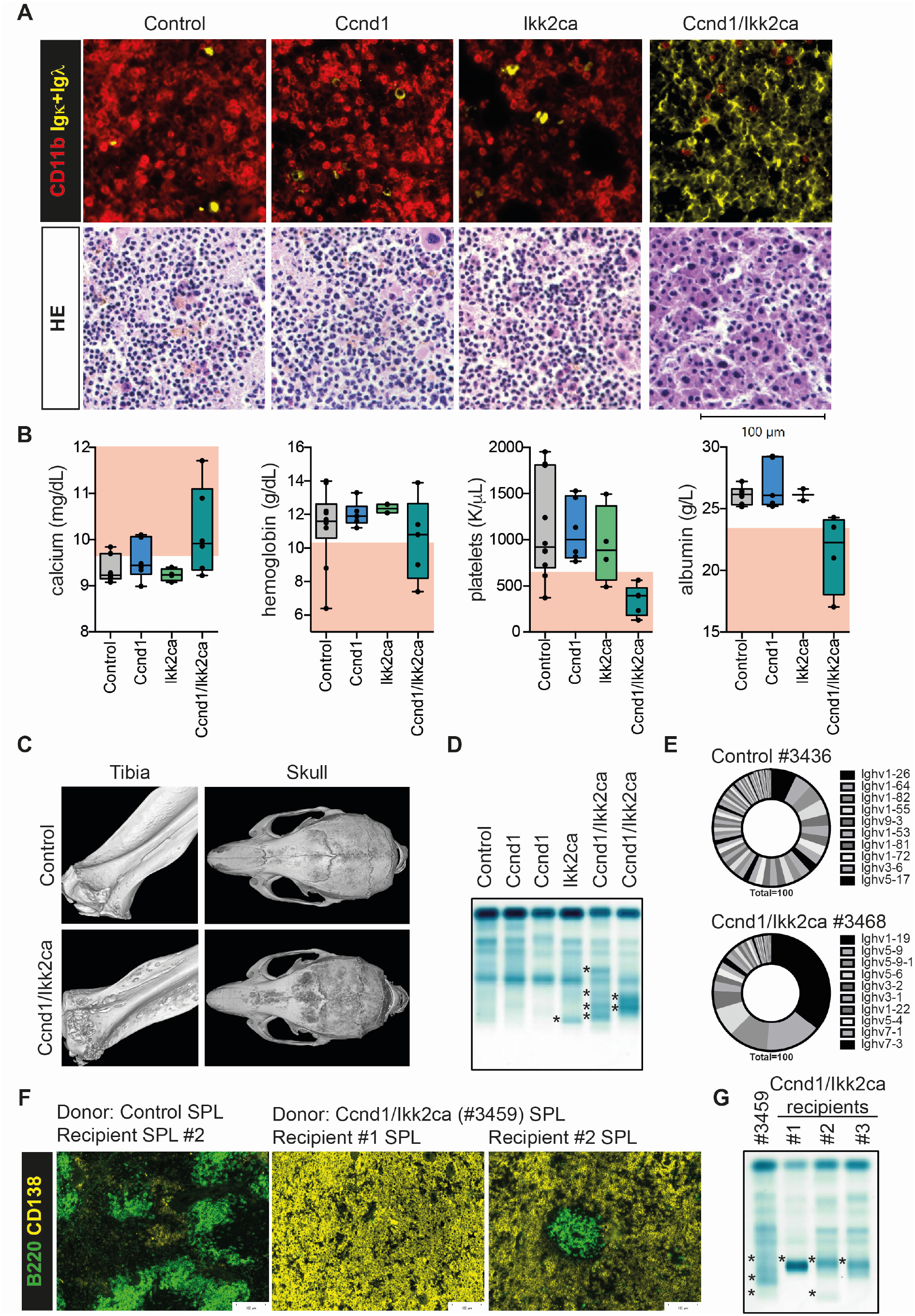
Ccnd1/Ikk2ca mice develop a plasma cell expansion and associated MM features with age. (A) Upper panel: representative immunofluorescence images of femur sections stained with α-CD11b (red, myeloid cells) and α-Igκ/α-Igλ (yellow, plasma cells). Lower panel: representative HE stained femur sections. (B) Graphs depicting serum calcium, hemoglobin, platelets and serum albumin values of Ccnd1 cohort mice (as in Fig. 1D). (C) 3D images reconstructed from µCT scans of tibiae and skulls. (D) Representative SPEP of Ccnd1 cohort mice showing the occurrence of M spikes (*). (E) Pie charts depicting the fraction of individual Ighv genes among all annotated Ighv gene reads of control and Ccnd1/Ikk2ca BM plasma cells. The 10 most dominantly expressed Ighv genes are listed (ordered clockwise). (F-G) Total spleen cell transfer of a control (as in Fig. 2D-F) or a Ccnd1/Ikk2ca (#3459) mouse into Rag2-Il2rg-recipients, followed by analysis after 31 weeks. (F) Representative images of recipient spleens stained with α-B220 (green, B cells) and α-CD138 (yellow, plasma cells). (G) SPEP of the donor (Ccnd1/Ikk2ca; #3459) and three recipient mice showing the occurrence of M proteins (*).

To link our model to human MM, we sorted BM plasma cells from aged control, single (Ccnd1 or MMSET or Ikk2ca) or double mutant (Ccnd1/Ikk2ca or MMSET/Ikk2ca) mice and performed RNA-Seq (Fig. 4A). Principal component (PC) analysis based on the top 500 genes showing the highest variability in expression across all samples accurately clustered the samples according to genotype (Fig. 4B). Double transgenic cells were clearly separated from control BM plasma cells by PC1, suggestive of an aberrant disease-related gene expression profile. Moreover, PC2 further divided double mutant cells into Ccnd1/Ikk2ca and MMSET/Ikk2ca samples reminiscent of sub-classification in human MM (Fig. 4B). Likewise, unsupervised hierarchical clustering positioned all double transgenic BM plasma cells into one branch, with clearly defined sub-branches for Ccnd1/Ikk2ca and MMSET/Ikk2ca samples (Fig. 4C). We then employed gene set enrichment analysis (GSEA) using the tmod algorithm ((*25*)) to determine whether (human) MM signature genes are enriched in double mutant plasma cells. To this end, we combined two published lists of genes differentially up- or down-regulated in MGUS/MM cells compared to normal human plasma cells into an MM signature_UP or MM signature_DOWN module, respectively ((*26*), (*27*)). Notably, the MM signature_UP was significantly enriched among genes up-regulated in Ccnd1/Ikk2ca or MMSET/Ikk2ca BM plasma cells in comparison to their respective controls, whereas the MM signature_DOWN was significantly enriched among the down-regulated genes (Fig. 4D-E), suggesting that the outgrowing mouse plasma cells share a transcriptional profile with human MGUS/MM cells. It should be noted that these signatures were also enriched in single mutant plasma cells (Fig. S15A-B), suggestive of a premalignant MGUS stage in these mice, which occasionally showed faint M proteins in the absence of disease (Fig. 2A, 3D, S13A, S4D, S10C). To investigate whether the transgenic plasma cells match to their respective human MM subgroup counterparts, we performed GSEA within the genes differentially expressed between MMSET/Ikk2ca versus Ccnd1/Ikk2ca and MMSET versus Ccnd1 samples, respectively (Fig. 4F-G, S15C), using previously described MM subgroup-specific gene sets ((*11*), (*12*), (*13*)). Strikingly, all three MMSET signatures were significantly enriched among MMSET-specific genes, linking MMSET/Ikk2ca and (also MMSET single mutant) plasma cells to the human t(4;14) subgroup (Fig. 4F, S15C). For Ccnd1-specific genes, only the comparison of MMSET versus Ccnd1 single mutant plasma cells resulted in a significant enrichment, namely the CD-2 signature ((*12*)), suggesting that transgenic Ccnd1 plasma cells show transcriptional features of the t(11;14) subgroup (Fig. 4G).

**Figure 4:**
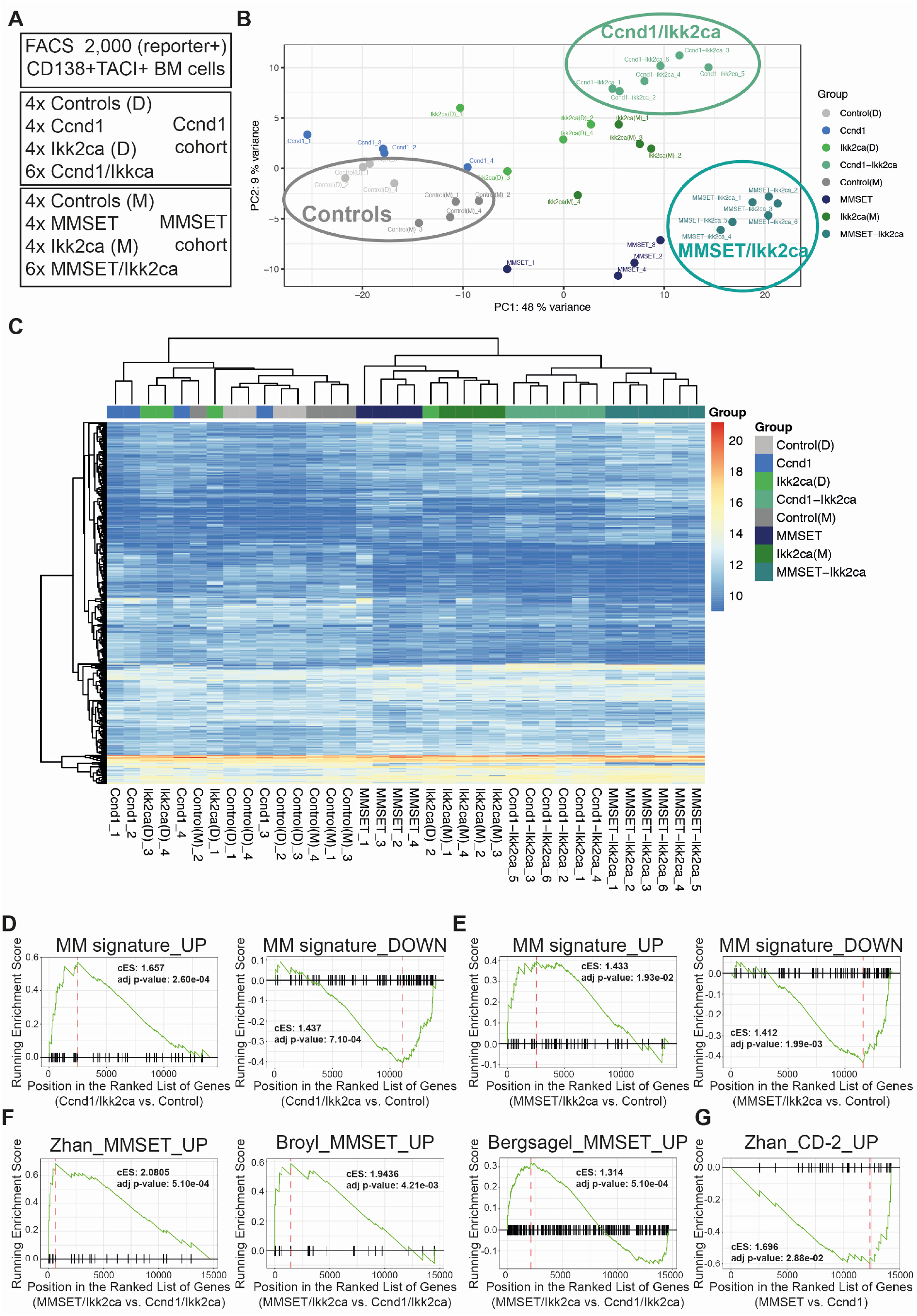
Transgenic mouse plasma cells express human MM signature genes. (A) Overview of RNA-sequencing samples. (B-C) Principal component analysis (B) and unsupervised hierarchical clustering (C) of control, single (Ccnd1, MMSET or Ikk2ca) and double (Ccnd1/Ikk2ca or MMSET/Ikk2ca) mutant BM plasma cells using the top 500 genes showing the highest variability in expression across all samples. (D-G) Graphical representations of gene set enrichment analysis results employing the tmod algorithm. (D-E) The x-axis shows the ranked gene list (ordered from most up-to most down-regulated) when comparing Ccnd1/Ikk2ca (D) or MMSET/Ikk2ca (E) to control BM plasma cells. The tested gene sets include genes differentially up-(MM signature_UP) or down-regulated (MM signature_DOWN) in human MGUS/MM cells versus normal human plasma cells ((*26*), (*27*)). (F) The x-axis shows the ranked gene list (ordered from most up-to most down-regulated) when comparing MMSET/Ikk2ca versus Ccnd1/Ikk2ca BM plasma cells. The tested gene sets include genes specifically up-regulated in the human t(4;14)/MMSET subgroup ((*11*), (*12*), (*13*)). (G) The x-axis shows the ranked gene list (ordered from most up-to most down-regulated) when comparing MMSET versus Ccnd1 BM plasma cells. The tested gene set includes genes specifically up-regulated in the human t(11;14)-associated CD-2 subgroup ((*12*)).

Taken together, our study reports novel genetically defined mouse models for distinct human MM subgroups. The GC B cell-specific co-activation of either Ccnd1 or MMSET, the two genes most frequently affected by a primary IGH translocation in human MM, with constitutive NFκB signaling as a secondary event, induced a clonally restricted plasma cell outgrowth, with most mice demonstrating > 60 % infiltration within the femur, reaching *per se* one of the newly-defined myeloma-defining events (MDE) ((*2*)). The significant but less prominent expansion of plasma cells measured by flow cytometry is likely a consequence of rapid cell death *ex vivo*, which is a known feature of human MM cells that strongly depend on their BM microenvironment ((*28*)). With respect to disease manifestations, the most apparent features of Ccnd1/Ikk2ca and MMSET/Ikk2ca mutant mice were hypoalbuminemia and thrombocytopenia – both of which have prognostic importance in human MM ((*29*)). In addition, classical CRAB criteria ((*2*)) were detected in a substantial proportion of double mutant mice validating the onset of an MDE also in mice with femur infiltration rates < 60 % (Table S1, S3). Gene expression profiling confirmed that the expanding mouse plasma cells were enriched for human MGUS/MM signature genes, further connecting our model to the human disease. Importantly, MMSET and Ccnd1 transgenic plasma cells could be linked to their respective human t(4;14) and t(11;14) subgroups: The t(4;14)/MMSET subgroup-specific signatures were significantly enriched within MMSET and MMSET/Ikk2ca BM plasma cells, whereas the t(11;14)-associated CD-2 signature was associated with Ccnd1 single mutant cells. The missing overlap of Ccnd1/Ikk2ca BM plasma cells with their human t(11;14) counterpart might be due to a dominant effect of Ikk2ca on the transcriptome, as indicated by the PCA that closely groups Ikk2ca and Ccnd1/Ikk2ca samples together (Fig. 4B).

Interestingly, only very few single mutant mice presented with a full disease phenotype, although some demonstrated serum M spikes along with a restricted BM plasma cell repertoire and a transcriptional profile of human MGUS/MM cells. This suggests that in contrast to double mutants, age-matched single mutants were either healthy or exhibited a premalignant stage, highlighting a) the relatively low oncogenic potential of the primary genetic aberrations on their own and b) the importance of synergistic effects of primary and secondary events for the development of overt MM. Despite the resemblance to human MM, some aspects of the disease are still not fully recapitulated. Important limitations include the substantial fraction of IgM isotype expression and the overall low degree of somatic hypermutation within the VDJ sequences of the expanding plasma cell clones (Tables S2, S4). These limitations are likely consequences of early Ikk2ca activation by the C*γ*1-cre transgene, since NFκB activation actively drives B cells out of the GC, promoting plasma cell differentiation ((*30*)). An expansion of IgM-expressing plasma cells is usually associated with Waldenström’s macroglobulinemia (WM), an indolent lymphoplasmacytic lymphoma ((*31*)). However, the occurrence of IgM MM has also been described, differing from WM in gene expression ((*32*)) and clinical manifestation ((*33*)). In line with this observation, Ccnd1/Ikk2ca and MMSET/Ikk2ca BM plasma cells were clearly distinguished from WM cells by GSEA using MM- and WM-specific gene sets ((*34*)) (Fig. S15D-E).

In summary, the MM models described here demonstrate a prominent outgrowth of selected plasma cell clones with onset of CRAB symptoms. The expanded plasma cells can be propagated *in vivo*, allowing subsequent functional analyses, e.g. genetically guided vulnerability and drug screening approaches or selective cell transfer experiments to address the long-standing question of the identity of MM initiating cells. Thus, our MM models represent a significant progress compared to existing transgenic models as they do not only recapitulate key features of the human disease, but also offer the opportunity to study the heterogeneity of human MM in a defined genetic system.

## Acknowledgements

We thank all members of the M. Janz & S. Mathas and K. Rajewsky groups for their feedback and discussion; K. Petsch, C. Salomon, J. Pempe and the MDC animal caretakers for their outstanding technical help; H.P. Rahn und K. Rautenberg for excellent FACS-related support; R. Kühn and the MDC Transgenic Core Facility for the generation of R26 Ccnd1^stopF^ and R26 MMSET^stopF^ alleles; W. Schneider for assessment of mouse kidney biopsies and S. Sauer and the MDC Genomics Facility for performing the RNA-sequencing run.

## Author’s Contributions

W.W., C.F.D., M.W., B.W. and B.W.W. performed experiments. W.W., E.B. and P.L. analysed data. H.N. performed µCT measurements and analyses. I.A. and A.R. provided pathological assessment. T.Y., A.H., D.B. and S.M. provided critical scientific, technical and experimental advice; W.W., K.R. and M.J. wrote the manuscript; and K.R. and M.J. conceived the study and supervised the project.

## Data and materials availability

The RNA-Seq data are in the process to be made available in the Gene Expression Omnibus (GEO) database (www.ncbi.nlm.nih.gov/geo; RRID:SCR_005012). The materials required to use any of the newly generated mouse strains will be available through K.R. and M.J. under a material transfer agreement from the MDC.

## Supplementary Information

Materials and Methods

Supplementary Figure Legends

Figures S1-S15

Supplementary Table Legends

Tables S1-S6

References (35-47)

