## Supplementary Information for "Preclinical models of human multiple myeloma subgroups"

*corresponding authors:

Supplementary Figure Legends

Figures S1-S15

Supplementary Table Legends

Tables S1-S6

References (35-47)

#### Materials and Methods

*Mice, Immunization, and Tumor Cohorts*

Cγ1-cre, R26 Ikk2ca$^{stopF}$ and R26 BFP$^{stopF}$ alleles have been described previously ((*23*), (*19*), (*35*)). A similar targeting strategy as for the Ikk2ca allele was applied to generate R26 MMSET$^{stopF}$ and R26 Ccnd1$^{stopF}$ alleles ((*19*)). In short, a construct encoding for either mouse Ccnd1 cDNA (NM_007631.3) or mouse MMSET-II cDNA (*Nsd2*; NM_001081102.2) preceded by a loxP-flanked STOP cassette was integrated into the mouse Rosa26 locus. Transgene transcription is controlled by a CAG promoter and its expression can be detected by the enhanced blue fluorescent protein (EBFP), which is placed under control of an internal ribosomal entry site (IRES) downstream of the respective cDNAs. The linearized targeting vector was transfected into the Artemis B6/3 C57BL/6 ES cell line and targeted clones were isolated using positive (NeoR) selection. Correct integration was verified by Southern blot of EcoRI- or PacI-digested genomic DNA from mouse ES cells and founder mouse tails using a Rosa26-specific probe (external Rosa probe A) ((*19*)) or Neo-specific (BamHI/PstI-digested fragment from NeoR) probe, respectively.

Mice were bred and maintained under specific pathogen-free conditions. Eight to 12 week old mice, indiscriminately of their sex, were immunized intraperitoneally with 100 µg alum-precipitated 4-hydroxy-3-nitrophenylacetyl-hapten conjugated to chicken gamma globulin (NP-CGG, Ratio10-19, LGC Biosearch Technologies Cat#N-5055B) and either analyzed after 14 days (Fig. S2) or monitored for tumor formation. Cohort mice included heterozygous control (Cγ1-cre only, R26 floxed or BFP reporter), single (Ccnd1, MMSET or Ikk2ca) and double (Ccnd1/Ikk2ca or MMSET/Ikk2ca) mutant mice. At the day of sacrifice, cohort mice were euthanized by cardiac puncture to obtain a sufficient blood sample for hematology and clinical chemistry. To this end, mice were first injected subcutaneously with 0.1 mg per kg body weight Buprenorphin (Buprenovet sine), followed by deep anesthesia with 4 Vol.-% isoflurane and subsequent puncture of the left ventricle. Hematologic and clinical serum parameters were measured on the ProCyte Dx (IDEXX) and AU480 Chemistry Analyser (Beckman Coulter), respectively. For the adoptive transfer, $1.15x10^6$ thawed total spleen cells of either BFP, MMSET/Ikk2ca or Ccnd1/Ikk2ca mice were injected intravenously into Rag2- ((*36*)) Il2rg- ((*37*)) recipients, which had been irradiated with 3 Gy one day before. Breeding, maintenance and experimental animal procedures were approved by the Landesamt für Gesundheit und Soziales Berlin (G0374/13, G0196/19, G0029/15, G0308/19).

*Flow cytometry and cell sorting*

Red blood cells were lysed with Gey's solution and single-cell suspensions from spleen as well as BM derived from either the long bones (femur, tibia, humerus) or the spine were stained with antibody conjugates (Table S5) in PBS, pH 7.2, supplemented with 3 % FCS and 1 mM EDTA. The samples were analyzed on an LSRFortessa (BD BioSciences) or sorted on a FACSAria (BD BioSciences). Plots were generated using FlowJo software (BD FlowJo, RRID:SCR_008520; v9.9.6). For the analysis of NIP-specific GC B cells, NIP-BSA-APC was used as previously described ((*38*)).

*B cell culture and TATCre treatment*

Splenic B cells were enriched by CD43 depletion with magnetic anti-mouse CD43 microbeads (Miltenyi Biotech, Cat#130-049-801) according to the manufacturer's instructions. To remove the STOP cassette, transgenic B cells were transduced *ex vivo* with in-house generated TATCre recombinase ((*39*)) and cultured in the presence of recombinant mouse IL-4 (25ng/mL, R&D, Cat#404-ML) and either LPS (20µg/mL, Sigma, Cat#L2880) or anti-mouse CD40 (1µg/mL, BioLegend, HM40-3, Cat#102908). For the confirmation of MMSET expression, B cells had to be differentiated into plasma cells using the 40LB feeder cell system and recombinant mouse IL-21 (10 ng/mL, Peprotech, Cat#210-21) ((*40*)).

*Real-time PCR and Western blotting*

BFP+ B or plasma cells were sorted, total RNA extracted using the RNeasy Mini kit (QIAGEN, Cat#74104) and cDNA synthesized using SuperScript Reverse Transcriptase (Invitrogen, Cat#18064022). The quantitative PCR was done with SYBR Green, followed by analysis with the StepOnePlus System (Applied Biosystems). Samples were assayed in triplicates, and mRNA abundance was normalized to that of *Hprt*. For Western blot, whole cell extracts of sorted BFP+ B or plasma cells were prepared as previously described ((*41*)). The primary antibodies used are listed in Table S5.

*Immunohistochemistry*

For decalcification, femurs were incubated for 24 h in 0.4 M EDTA pH 7.4, followed by 24 h in 30 % sucrose solution (in PBS) under constant rotation at 4°C. Spleen sections and decalcified bones were embedded in Tissue-Tek O.C.T. Compound (Sakura Cat#4583), stored at -80°C and cryosectioned (7µm thickness). For immunofluorescence staining, slides were fixed in 100 % acetone, blocked with 3 % BSA/PBS and stained with DAPI (eBioScience Cat# D1306) and the antibody conjugates listed in Table S5. For HE staining, frozen sections from the femur and spleen were stained with hematoxylin and eosin (HE) according to the following protocol: the slides were immersed for 15 min in hemalaun solution followed by a 5 min rinse in tap water and a 5 min incubation in eosin solution. After short (5 sec) rinses in tap water followed by 100% ethanol and xylene, the slides were mounted in a film coverslipper (Tissue-Tek-Film® Automated Coverslipper, Sakura Finetek USA) and evaluated in a blinded fashion, i.e. without knowledge of the genotypes. Immunoflourescence and HE stained sections were imaged with a BZ-9000 microscope (Keyence).

*Enzyme-Linked Immuno Assays, Serum Protein Electrophoresis and Immunofixation*

Enzyme-linked immunosorbent assays (ELISAs), enzyme-linked immuno spot (ELISPOT) assays and serum protein electrophoresis were performed as described with the minor change that for the ELISA of cohort mouse samples plates were coated with 1 µg/mL anti-IgM, anti-IgG or anti-IgA instead of anti-light chain antibodies ((*38*)). Immunofixation was done on 1:6 (for IgG) or 1:3 (for all other Ig isotypes) diluted serum samples according to the "Hydragel 4 IF Immunofixation" kit (Sebia, Cat# 4804) using mouse antisera listed in Table S5.

*VDJ-PCR*

Genomic DNA was prepared from sorted BFP+GFP+ CD138+TACI+ long bone-derived BM plasma cells (NucleoSpin Tissue, Macherey-Nagel, Cat#740952). Ighv gene rearrangements were PCR amplified using the KOD HotStart DNA polymerase (Novagen, Cat#71086) and forward primers specific for VHA, VHB, VHC, VHD, VHE, VHF, VHG ((*24*)) as well as a reverse primer in the $J_H4$ intron ((*30*)). Prominent fragments were cloned, sequenced and aligned against the NCBI database IgBLAST (http://www.ncbi.nlm.nih.gov/igblast/; RRID:SCR_002873) to determine $V_HD_HJ_H$ gene segment usage.

*RNA-sequencing*

2,000 (reporter+) CD138+TACI+ long bone-derived BM plasma cells from 8x control, 4x Ccnd1, 4x MMSET, 8x Ikk2ca, 6x Ccnd1/Ikk2ca and 6x MMSET/IKk2ca mice were separately sorted into 75 µL RLT buffer and stored at -80°C. Total RNA was extracted using the RNeasy Micro Kit (QIAGEN, Cat#74004) and subsequently used to generate cDNA libraries according to a low-input protocol based on sequence-independent full-length cDNA amplification ((*42*)). Libraries were sequenced at the Genomics facility of the Max-Delbrück-Centrum on a NovaSeq6000 instrument as single 150 bp reads with a depth of approximately 100 million reads per sample. RNA-Seq data are in the process of being deposited at the GEO (Gene Expression Omnibus; https://www.ncbi.nlm.nih.gov/geo/; RRID:SCR_005012) repository.

*Bioinformatic analysis: GSEA and Ighv gene usage*

The 36 sequenced libraries contained between 71.8 & 130.4 million reads (median 86.5 million reads). The reads were mapped on the GRCm38 genome (GENCODE m23 annotations) using salmon ((*43*)) version 1.3.0 in selective alignment mapping mode. The mapping rate was between 61.0 % and 68.9 % (median 64.7 %). Because of their high abundance, genes and pseudogenes from the IMGT repertoire (immunoglobulins, J chain gene and T-cell receptors) were excluded from the gene expression analysis. The gene expression was computed from the mapping on transcripts using the tximport bioconductor package ((*44*)). DESeq2 ((*45*)) was used to compute differential expression over all samples together. Normalised expression levels were obtained by variance-stabilisation transformation. For the gene set enrichment analysis, an overview of the tested gene sets can be found in Table S6. The tested modules were derived from Tables 4 and 5 of (*26*), Table S4 from (*27*), Tables 3 and 4 from (*34*) and all supplementary tables containing MM subgroup-specific gene lists from (*11*), (*12*) and (*13*). To compute the gene sets' statistical significance, the CERNO test from R package tmod ((*25*), (*46*)) was used. For the CERNO test, mouse genes were ranked according to the value of the differential expression statistic provided by the "results" function of DESeq2. For every gene set, the test was run in both directions (from up- to down-regulation, and from down- to up-regulation). The clusterProfiler biocondutor package ((*47*)) was used to produce the gene set enrichment plots. Gene symbols required by the functional analysis were obtained directly from the GENCODE annotations.

For the Ighv (immunoglobulin heavy chain variable region) gene usage analysis, due to the 129 genetic background of the Cγ1-cre allele ((*23*)), counts for the Ighv genes were obtained by mapping reads on the GRCm38 genome, augmented with Ighv transcripts from the 129S1/SvImJ genome (taken from the ENSEMBL 103 release). The mapping was done using salmon. Then, for each sample the counts aligning to an individual Ighv gene were normalized to all Ighv gene counts (i.e. by setting the sum of all Ighv gene reads to 100 %) and plotted as a pie chart.

*μCT analysis*

Mouse skulls and tibiae were scanned with a SkyScan 1276 scanner (Bruker, Belgium) using the vendor's software for image acquisition (v.1.4.0.0), the step-and-shoot mode, the source current of 200 μA and 360° acquisition. Tibiae were scanned using the source voltage of 55 kV, Al 0.5 mm filter, an exposure time of 640 ms, 0.1° rotation step, and 3 frame averages. The pixel size was 5.0 μm. The skulls were scanned with the source voltage of 100 kV, Cu 0.25 mm filter, exposure time of 645 ms, 0.2° rotation step, and 3 frame averages. The pixel size was 10.65 μm. The flat field correction was applied for all acquisitions. Image reconstruction was performed with NRecon (v.1.7.3.1, Bruker, Belgium), using the beam-hardening correction of 26 % and the ring-artefact correction of 6. No smoothing was applied. All μCT images were analyzed qualitatively using CTVox (v.3.3.1.0, Bruker, Belgium). Severity grading of tibiae included "normal" for tibiae with < 3 clear-rimmed lesions and "pathologic" for tibiae with ≥ 3 clear-rimmed lesions. Of note, some tibiae demonstrated small (likely age-related) spots of qualitatively lower bone density, which appearance differed from the defined lesions and were excluded from the severity grading. Skull images were graded as "normal" as long as areas of qualitatively lower bone density were absent or only locally restricted and "pathologic" once these areas were diffuse and/or accompanied with lesions.

*Data presentation and Statistical Analysis*

Single dots represent number of biological replicates from independent mice. Unless noted otherwise, bars or lines represent median values. Prism software (GraphPad

Prism, RRID: SCR_002798) version 8 was used for pair-wise comparisons between control or single mutant and respective double mutant samples using non-parametric, unpaired, two-tailed Mann-Whitney U test. Asterisks indicate statistical significance for p-values < 0.05 (single), < 0.01 (double), < 0.001 (triple) and < 0.0001 (quadruple). For representation of blood parameters, box-and-whisker graphs are shown with boxes extending from the 25$^{th}$ to 75$^{th}$ percentile with a line at the median and whiskers plotted down to the minimum and up to the maximum value.

**Supplementary Figure Legends**

**Figure S1: Gene targeting strategy and validation of transgene expression.** (A) Scheme of the wildtype (WT) and targeted (T) Rosa26 locus. Ex = exon; CAG = CAG promoter; Neo = Neomycin resistance gene; STOP = STOP cassette; PRIM = primary candidate, i.e. Ccnd1 or MMSET; I = IRES (internal ribosomal entry site); BFP = blue fluorescent protein; R4A and neo denote Southern blot probes. (B) Southern blot for R4A and neo probes on EcoRI- or PacI-digested tail genomic DNA of the indicated mouse genotypes. Note: The *MMSET* isoform RE-IIBP was also targeted but not used for this study. (C) Experimental scheme to check for transgene expression in stimulated B cells *in vitro*. (D) Quantitative PCR of BFP+ B cells sorted two days after TATCre treatment. The primers detect transgenic *Ccnd1* (Ccnd1-tg), transgenic *MMSET* (MMSET-tg) and *Hprt* (control) mRNA expression, respectively. (E) Immunoblot for Ccnd1 on whole cell lysates of BFP+ B cells sorted at day 2. C57BL/6 B cells retrovirally transduced with empty vector (MIG) or MIG-Ccnd1 as well as human MM cell lines served as controls. (F) Experimental scheme for the *in vitro* generation of plasma cells using the Kitamura 40LB ((*40*)) system. (G) Gating strategy for BFP+ plasma cells. Contour plot depicts plasma cells as B220lowCD138+ cells which were then separated into BFP+ and BFP- cells and sorted. (H) Immunoblot for MMSET on whole cell lysates of BFP+ and BFP- plasma cells derived from 40LB-cultured R26 MMSET$^{stopF}$ cells. ß-actin (Actb) served as loading control.

**Figure S2: Analysis of the immune response to NP-CGG.** (A) Immunization scheme. Control (BFP), single mutant (Ccnd1 or MMSET or Ikk2ca) and double mutant (Ccnd1/Ikk2ca or MMSET/Ikk2ca) mice were immunized once with NP-CGG and analyzed at day 14. Right: Gating strategy for antigen (NIP)-specific GC B cells (pre-gate: living B220+CD19+ B cells). (B) Absolute numbers of splenic NIP-specific IgG1+ GC B cells from mice with the indicated genotypes 14 days after immunization; each dot represents one mouse. (C) NP-specific serum IgG1 titers measured by ELISA. (D-E) Numbers of NP-specific IgM- or IgG1-producing cells within 0.16E06 splenocytes (left) or 0.8E06 BM cells (right) of Ccnd1 (D) or MMSET (E) cohort mice. Statistics: Mann-Whitney-test; *p<0.5; **p<0.01; ***p<0.001; ****p<0.0001

**Figure S3: Characterization of the plasma cell expansion in aged MMSET/Ikk2ca mice.** (A) Spleen weight and (B) total cell numbers of the spleen and BM (isolated from one tibia, one femur and one humerus) of mice with the indicated genotypes. (C) Representative flow cytometry contour plots of BM isolated from mice with the indicated genotypes depicting CD138+TACI+ plasma cells (upper panel) and reporter expression within this population (lower panel). (D) Percentage of CD138+TACI+ plasma cells within the spleen, long bone-derived BM and spine-derived BM. (E) Absolute numbers of CD138+TACI+ plasma cells within the spleen and long bone-derived BM. (F) Reporter (BFP and/or GFP) expression within CD138+TACI+ plasma cells of the spleen, long bone-derived BM and spine-derived BM. (G) Representative immunofluorescence images of spleen sections stained with $\alpha$-CD19 (green, B cells) and $\alpha$-CD138 (yellow, plasma cells). (H) ELISA of total serum IgM, IgG and IgA titers. (I) ELISPOT analysis of immunoglobulin isotypes (combination of $\alpha$-IgM, $\alpha$-IgG1, $\alpha$-IgG2a/b/c, $\alpha$-IgG3, $\alpha$-IgA) on thawed long bone-derived total BM cells. Statistics: Mann-Whitney-test; *p<0.5; **p<0.01; ***p<0.001; ****p<0.0001

**Figure S4: MM-associated pathologies of MMSET cohort mice.**

(A-B) Representative 3D images reconstructed from µCT scans of tibiae (A) and skulls (B). Tibiae are shown in frontal view (A; left) and as zoomed-in image (A; right) to demonstrate that lesions destroyed the cortex. (C) Graph depicting red blood cell counts of cohort mice with the indicated genotypes at the day of sacrifice. The lower pathologic range of red blood cells (< 6.8 M/µL) is marked in red. (D) Heatmap summarizing the phenotyping of MMSET cohort mice. Columns represent selected MM-associated parameters; from left to right: ≥ 10 % percent of BM plasma cells (BMPC) measured by flow cytometry (FC); ≥ 10 % percent BMPC in the femur biopsy (histology); serum calcium, hemoglobin, albumin, red blood cells (RBC), platelets (PLT), tibia lesions and skull osteopenia. For blood parameters the pathologic range is indicated in brackets (reference values: Jackson Laboratory; 78 weeks old C57BL/6J; *the lower limit for albumin had to be adjusted for our cohort to discern healthy and diseased mice for animals > 78 weeks). Each row represents an individual mouse, ordered by genotype. White marks indicate that the parameter is within the normal range, red marks indicate that the parameter is within the defined pathologic range and grey marks indicate that the parameter was not measured. For the µCT analysis, red marks indicate mice with ≥ 3 lesions within the tibiae and diffuse osteopenia of the skull.

**Figure S5: Ighv gene usage of control, MMSET, Ikk2ca and MMSET/Ikk2ca BM plasma cells.** 2,000 long bone-derived transgenic (reporter+) CD138+TACI+ BM plasma cells of aged cohort mice were sorted and analyzed by RNA-sequencing. (A) Pie charts depict the fraction of individual Ighv genes among all annotated Ighv gene reads for cohort mice of the indicated genotypes. Shown are the results for all animals of the entire cohort analyzed by RNA-seq. The 10 most dominantly expressed Ighv genes are listed. Pie charts need to be read clockwise. Note: only 1,000 plasma cells could be sorted from MMSET mouse #3846. (B) Graph illustrating the fraction of the most dominant Ighv gene per mouse of the indicated genotypes.

**Figure S6: VDJ-PCR analysis of MMSET/Ikk2ca cohort mice.** Genomic DNA was isolated from sorted reporter+ CD138+TACI+ BM plasma cells (left panel), a femur slice (middle panel) or a spleen slice (right panel) and subjected to VDJ-PCR analysis covering the Ighv gene families (J558 (VHA), Q52 (VHB), 36-60 (VHC), X24 (VHD), 7183 (VHE), J606 and S107 (VHF) and GAM3 (VHG) ((*24*)). Orange boxes mark shared clones between sorted cells and either femur or spleen sections.

**Figure S7: Immunoglobulin isotype expression of MMSET/Ikk2ca plasma cells.** (A) SPEP coupled to immunofixation of the six MMSET/Ikk2ca mice whose BM plasma cells were also used for RNA-sequencing showing the Ig heavy chain and Ig light chain isotypes of the detected M proteins. Corresponding Ig light and heavy chains are marked with the same symbol; abbreviations: M - IgM, G - IgG, K - kappa light chain, L - lambda light chain. (B) Pie charts depict the fraction of individual Igh isotypes among all annotated Igh isotype gene reads determined by RNA-sequencing of sorted BFP+GFP+ CD138+TACI+ MMSET/Ikk2ca BM plasma cells.

**Figure S8: Analysis of MMSET/Ikk2ca (#3939) spleen cell recipient mice.** (A) Representative contour plots depicting CD138+TACI+ plasma cells and the reporter expression within this population. Upper panel: MMSET/Ikk2ca donor spleen; lower panel: representative recipient spleen and long bone-derived BM. (B) Shown are absolute numbers of CD138+TACI+ plasma cells in the spleen and long bone-derived BM (upper panel) and the enrichment of plasma cells in recipient mice compared to the number of transplanted plasma cells ($0.47 \times 10^6$ transferred plasma cells/recipient; lower panel). (C) Immunofluorescence images of spleen sections stained with $\alpha$-B220 (green, B cells) and $\alpha$-CD138 (yellow, plasma cells) of all recipient mice that received either control (BFP) or MMSET/Ikk2ca (#3939) splenocytes.

**Figure S9: Characterization of the plasma cell expansion in aged Ccnd1/Ikk2ca mice.** (A) Spleen weight and (B) total cell numbers of the spleen and BM (isolated from one tibia, one femur and one humerus) of mice with the indicated genotypes. (C) Representative contour plots of the long bone-derived BM depicting CD138+TACI+ plasma cells (upper panel) and reporter expression within this population (lower panel). (D) Percentage of CD138+TACI+ plasma cells within the spleen, long bone-derived BM and spine-derived BM. (E) Absolute numbers of CD138+TACI+ plasma cells within the spleen and long bone-derived BM. (F) Relative reporter (BFP and/or GFP) expression within CD138+TACI+ plasma cells of the spleen, long bone-derived BM and spine-derived BM. (G) Representative immunofluorescence images of spleen sections stained with $\alpha$-CD19 (green, B cells) and $\alpha$-CD138 (yellow, plasma cells). (H) ELISA of total serum IgM, IgG and IgA titers. (I) ELISPOT analysis of immunoglobulin isotypes (combination of $\alpha$-IgM, $\alpha$-IgG1, $\alpha$-IgG2a/b/c, $\alpha$-IgG3, $\alpha$-IgA) on thawed total BM cells. Statistics: Mann-Whitney-test; *$p<0.5$; **$p<0.01$; ***$p<0.001$; ****$p<0.0001$

**Figure S10: Analysis of MM-associated pathologies of Ccnd1 cohort mice.** (A) Representative 3D images reconstructed from µCT scans of tibiae (A) shown in frontal view (A; left) and as zoomed-in image (A; right) to demonstrate that lesions destroyed the cortex. (B) Graph depicting red blood cell counts of cohort mice at the day of sacrifice. The lower pathologic range of red blood cells (< 6.8 M/µL) is marked in red. (C) Heatmap summarizing the phenotyping of Ccnd1 cohort mice. Columns represent selected MM-associated parameters; from left to right: ≥ 10 % of BM plasma cells (BMPC) measured by flow cytometry (FC); ≥ 10 % of BMPC in the femur biopsy (histology); serum calcium, hemoglobin, albumin, red blood cells (RBC), platelets (PLT), tibia lesions and skull osteopenia. For blood parameters the pathologic range is indicated in brackets (see Fig. S4D). Each row represents an individual mouse, ordered by genotype. White marks indicate that the parameter is within the normal range, red marks indicate that the parameter is within the defined pathologic range and grey marks indicate that the parameter was not determined. For the µCT analysis, red marks indicate mice that demonstrated > 3 lesions within the tibiae and diffuse osteopenia of the skull.

**Figure S11: Ighv gene usage of control, Ccnd1, Ikk2ca and Ccnd1/Ikk2ca BM plasma cells.** 2,000 long bone-derived transgenic (reporter+) CD138+TACI+ BM plasma cells of aged cohort mice were sorted and analyzed by RNA-sequencing. (A) Pie charts show the fraction of individual Ighv genes among all annotated Ighv gene reads for all cohort mice of the indicated genotypes. The 10 most dominantly expressed Ighv genes are listed. Pie charts need to be read clock-wise. Note: Ikk2ca mouse #3299 presented with a monoclonal B cell tumor. (B) Graph illustrating the fraction of the most dominant Ighv gene per mouse within the groups of control, single mutant and double mutant mice.

**Figure S12: VDJ-PCR analysis of Ccnd1/Ikk2ca cohort mice.** Genomic DNA was isolated from sorted BFP+GFP+ CD138+TACI+ BM plasma cells (left panel), a femur slice (middle panel) or a spleen slice (right panel) and subjected to VDJ-PCR analysis covering the Ighv gene families (J558 (VHA), Q52 (VHB), 36-60 (VHC), X24 (VHD), 7183 (VHE), J606 and S107 (VHF) and GAM3 (VHG) ((*24*)). Orange boxes mark shared clones between sorted BM plasma cells and either femur or spleen sections.

**Figure S13: Immunoglobulin isotype expression of Ccnd1/Ikk2ca plasma cells.** (A) SPEP of the remaining control, Ccnd1, Ikk2ca and Ccnd1/Ikk2ca cohort mice demonstrating the emergence of M proteins (marked with an asterisk). (B) SPEP coupled to immunofixation of the six Ccnd1/Ikk2ca mice whose BM plasma cells were also used for the RNA-sequencing showing the Ig heavy chain and Ig light chain isotypes of the detected M proteins. Corresponding Ig light and heavy chains are marked with the same symbol. (C) Pie charts depicting the fraction of individual Igh isotypes among all annotated Igh isotype gene reads determined by RNA-sequencing of sorted BFP+GFP+ CD138+TACI+ Ccnd1/Ikk2ca BM plasma cells.

**Figure S14: Analysis of Ccnd1/Ikk2ca (#3459) spleen cell recipient mice.** (A) Representative contour plots depicting CD138+TACI+ plasma cells and the reporter expression within this population. Upper panel: Ccnd1/Ikk2ca donor spleen; lower panel: representative recipient spleen and long bone-derived BM. (B) Shown are absolute cell numbers of CD138+TACI+ plasma cells in the spleen and long bone-derived BM (upper panel) and the enrichment over the input amount of plasma cells ($0.2x10^6$ transplanted plasma cells/recipient; lower panel). (C) SPEP coupled to immunofixation to determine the Ig heavy chain and Ig light chain isotypes of the detected M proteins in the Ccnd1/Ikk2ca donor mouse (#3459) and its four recipients.

**Figure S15: GSEA analyses of human MM and Waldenström's Macroglobulinemia (WM) signature genes in transgenic mouse plasma cells (extension to Fig. 4).** Graphical representation of gene set enrichment analyses employing the tmod algorithm (*25*). (A-B) The x-axis shows the ranked gene list (ordered from most up- to most down-regulated) when comparing MMSET (A) or Ikk2ca (B) to control mouse BM plasma cells. The tested gene sets include genes differentially up- (MM signature_UP) or down-regulated (MM signature_DOWN) in human MGUS/MM cells versus normal human plasma cells ((*26*), ((*27*)). (C) The x-axis shows the ranked gene list (ordered from most up- to most down-regulated) when comparing MMSET and Ccnd1 BM plasma cells. The tested gene sets includes genes specifically up-regulated in the t(4;14)/MMSET subgroup of MM patients ((*11*), (*12*)). (D-E) The x-axis shows the ranked gene list (ordered from most up- to most down-regulated) when comparing Ccnd1/Ikk2ca (D) or MMSET/Ikk2ca (E) to control mouse BM plasma cells. The tested gene sets include genes specifically up-regulated in human MM (Gutierrez_MM) or human WM (Gutierrez_WM) cells versus normal human plasma cells ((*34*)).

### Figure S1

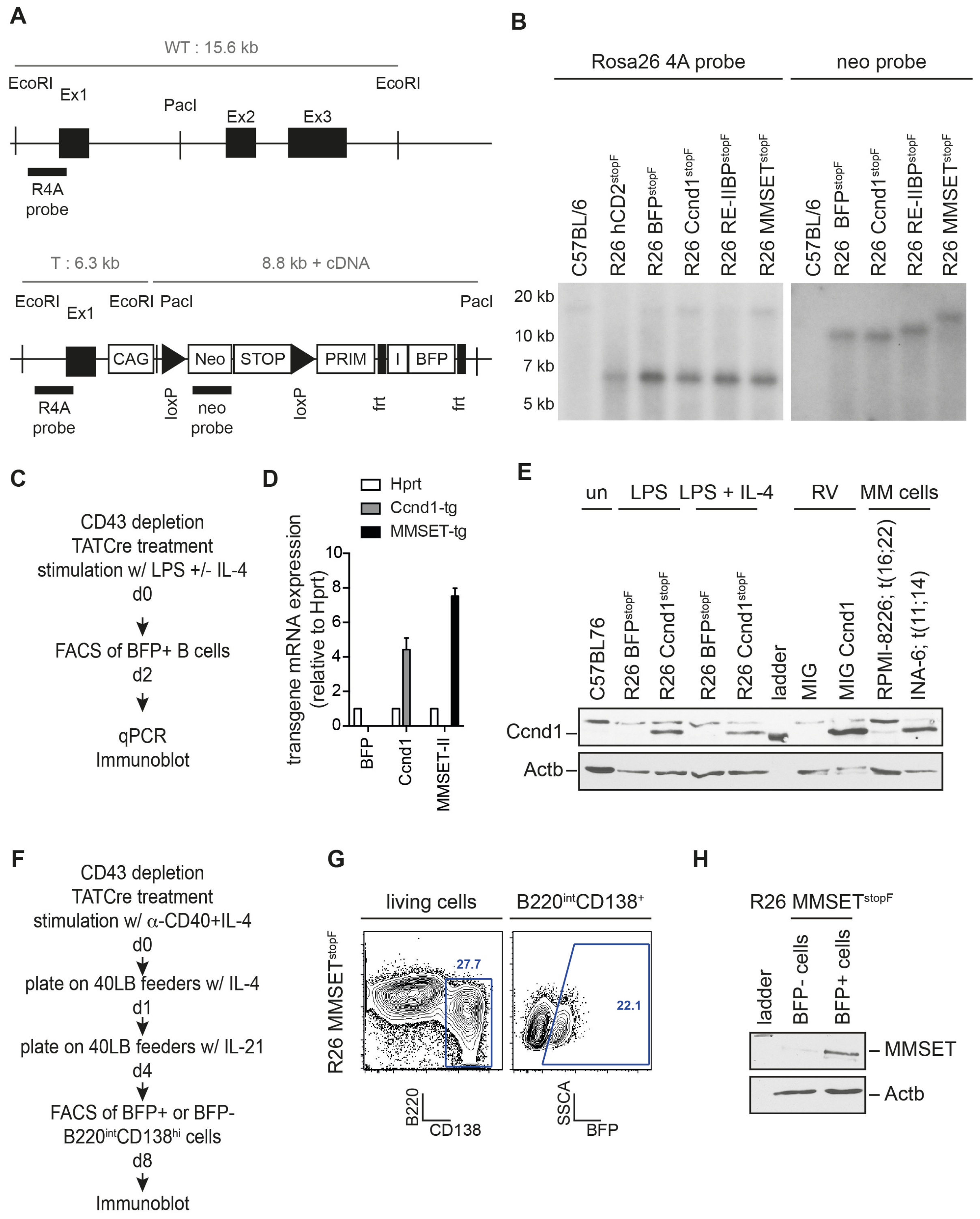

A
WT : 15.6 kb
EcoRI
Ex1
PacI
Ex2
Ex3
EcoRI
R4A probe
T : 6.3 kb
8.8 kb + cDNA
EcoRI
Ex1
EcoRI PacI
PacI
CAG
Neo
STOP
PRIM
I
BFP
R4A probe
loxP
neo probe
loxP
frt
frt
B
Rosa26 4A probe
neo probe
C57BL/6
R26 hCD2stopF
R26 BFPstopF
R26 Ccnd1stopF
R26 RE-IIBPstopF
R26 MMSETstopF
C57BL/6
R26 BFPstopF
R26 Ccnd1stopF
R26 RE-IIBPstopF
R26 MMSETstopF
20 kb
10 kb
7 kb
5 kb
C
CD43 depletion
TATCre treatment
stimulation w/ LPS +/- IL-4
d0
FACS of BFP+ B cells
d2
qPCR
Immunoblot
D
Hprt
Ccnd1-tg
MMSET-tg
transgene mRNA expression (relative to Hprt)
10
8
6
4
2
0
BFP
Ccnd1
MMSET-II
E
un
LPS
LPS + IL-4
RV
MM cells
C57BL76
R26 BFPstopF
R26 Ccnd1stopF
R26 BFPstopF
R26 Ccnd1stopF
ladder
MIG
MIG Ccnd1
RPMI-8226; t(16;22)
INA-6; t(11;14)
Ccnd1
Actb
F
CD43 depletion
TATCre treatment
stimulation w/ α-CD40+IL-4
d0
plate on 40LB feeders w/ IL-4
d1
plate on 40LB feeders w/ IL-21
d4
FACS of BFP+ or BFP-
B220intCD138hi cells
d8
Immunoblot
G
living cells
B220intCD138+
R26 MMSETstopF
27.7
22.1
B220
CD138
SSCA
BFP
H
R26 MMSETstopF
ladder
BFP- cells
BFP+ cells
MMSET
Actb

#### Figure S2

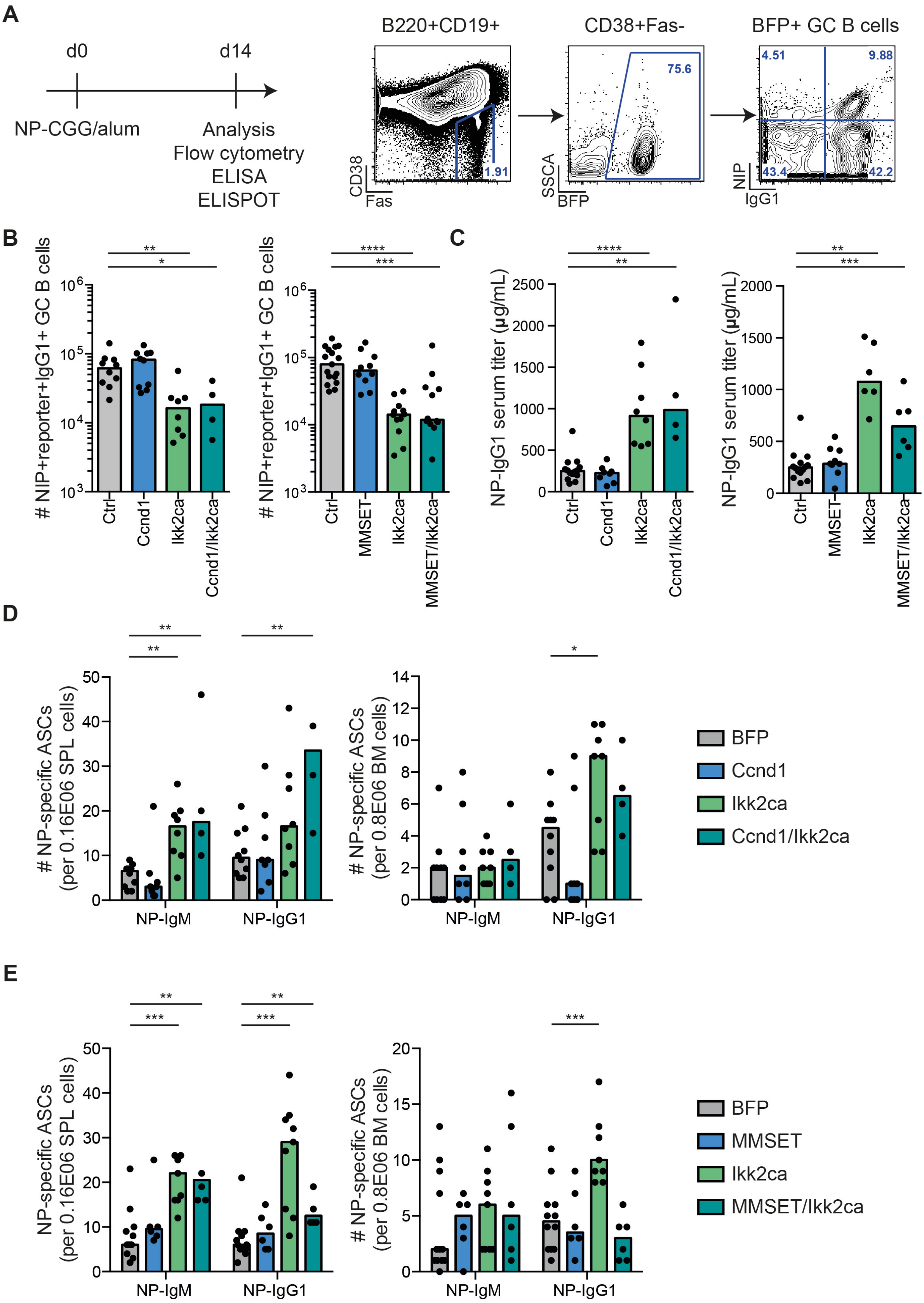
A
d0
d14
NP-CGG/alum
Analysis
Flow cytometry
ELISA
ELISPOT
B220+CD19+
CD38+Fas-
BFP+ GC B cells
CD38
Fas
1.91
SSCA
BFP
75.6
NIP
IgG1
4.51
9.88
43.4
42.2
B
### NIP+reporter+IgG1+ GC B cells
Ctrl
Ccnd1
Ikk2ca
Ccnd1/Ikk2ca
MMSET
MMSET/Ikk2ca
C
NP-IgG1 serum titer (µg/mL)
D
### NP-specific ASCs
(per 0.16E06 SPL cells)
(per 0.8E06 BM cells)
NP-IgM
NP-IgG1
BFP
Ccnd1
Ikk2ca
Ccnd1/Ikk2ca
E
NP-specific ASCs
(per 0.16E06 SPL cells)
### NP-specific ASCs
(per 0.8E06 BM cells)
BFP
MMSET
Ikk2ca
MMSET/Ikk2ca

**Figure S3**

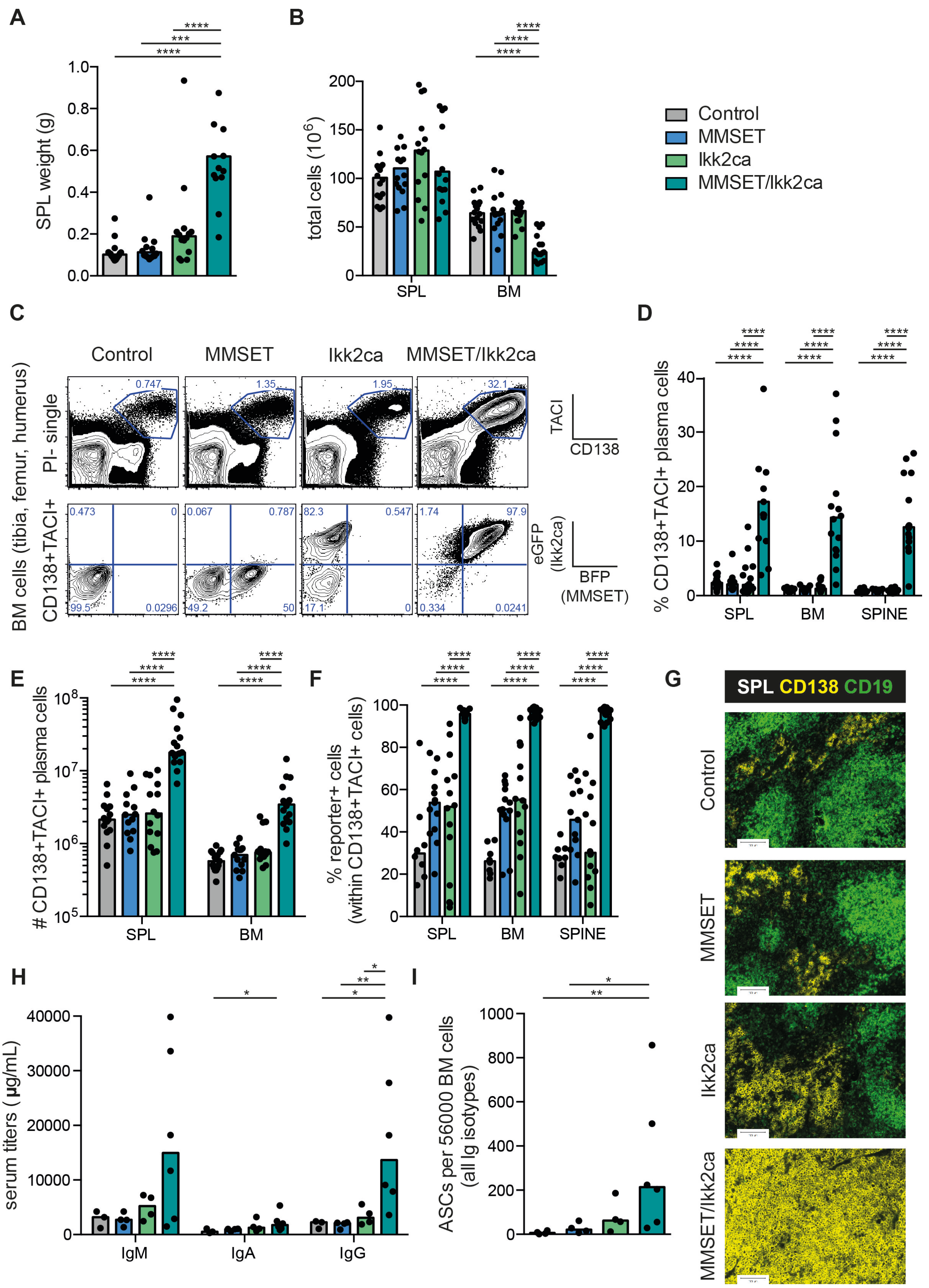
A
SPL weight (g)
B
total cells (10^6)
SPL
BM
Control
MMSET
Ikk2ca
MMSET/Ikk2ca
C
Control
MMSET
Ikk2ca
MMSET/Ikk2ca
BM cells (tibia, femur, humerus)
PI- single
CD138+TACI+
TACI
CD138
eGFP (Ikk2ca)
BFP (MMSET)
D
% CD138+TACI+ plasma cells
SPL
BM
SPINE
E
### CD138+TACI+ plasma cells
F
% reporter+ cells (within CD138+TACI+ cells)
G
SPL CD138 CD19
Control
MMSET
Ikk2ca
MMSET/Ikk2ca
H
serum titers (µg/mL)
IgM
IgA
IgG
I
ASCs per 56000 BM cells (all Ig isotypes)

#### Figure S4

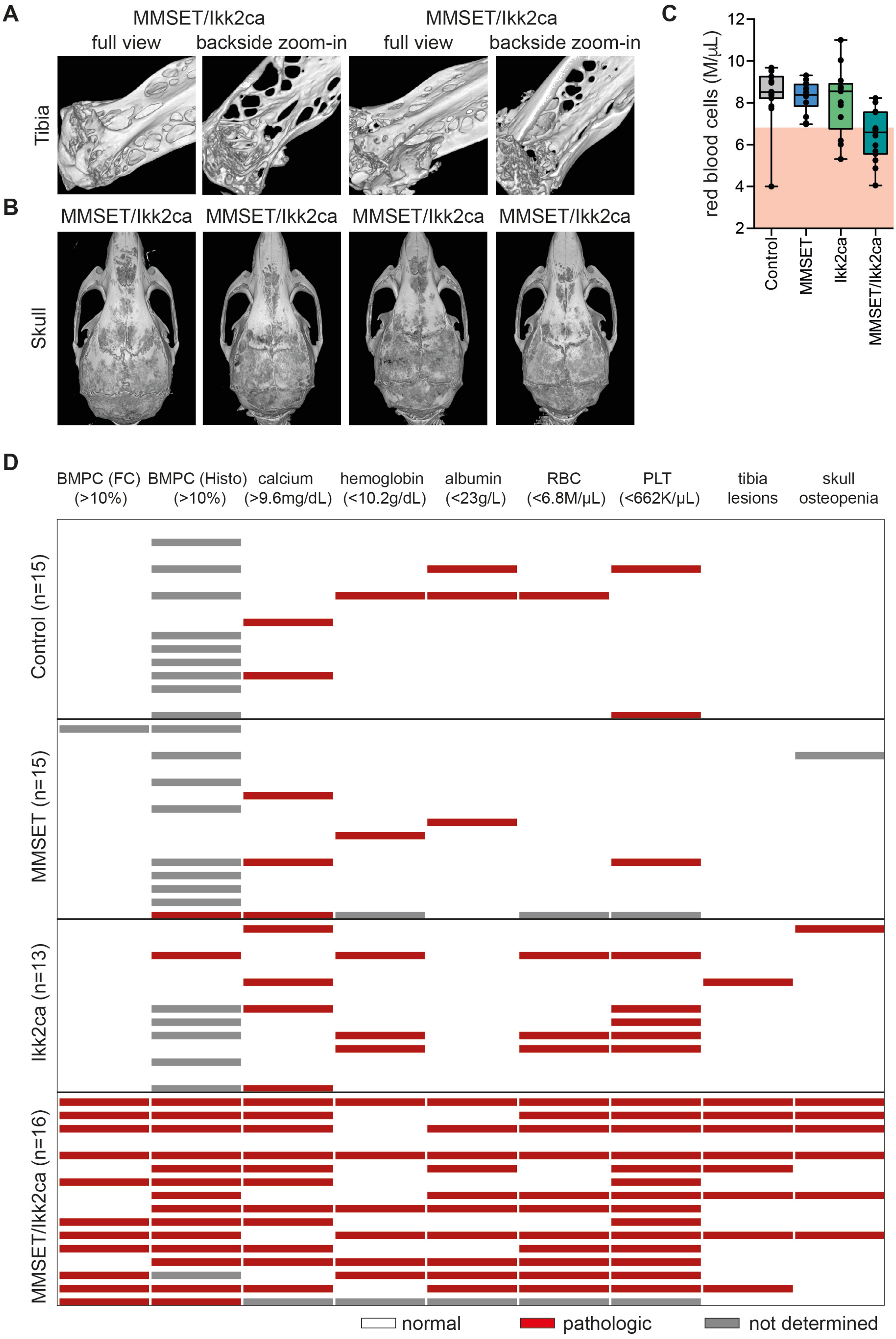
A
MMSET/Ikk2ca
full view
backside zoom-in
MMSET/Ikk2ca
full view
backside zoom-in
Tibia
B
MMSET/Ikk2ca
MMSET/Ikk2ca
MMSET/Ikk2ca
MMSET/Ikk2ca
Skull
C
red blood cells (M/μL)
12
10
8
6
4
2
Control
MMSET
Ikk2ca
MMSET/Ikk2ca
D
BMPC (FC) (>10%)
BMPC (Histo) (>10%)
calcium (>9.6mg/dL)
hemoglobin (<10.2g/dL)
albumin (<23g/L)
RBC (<6.8M/μL)
PLT (<662K/μL)
tibia lesions
skull osteopenia
Control (n=15)
MMSET (n=15)
Ikk2ca (n=13)
MMSET/Ikk2ca (n=16)
normal
pathologic
not determined

### Figure S5

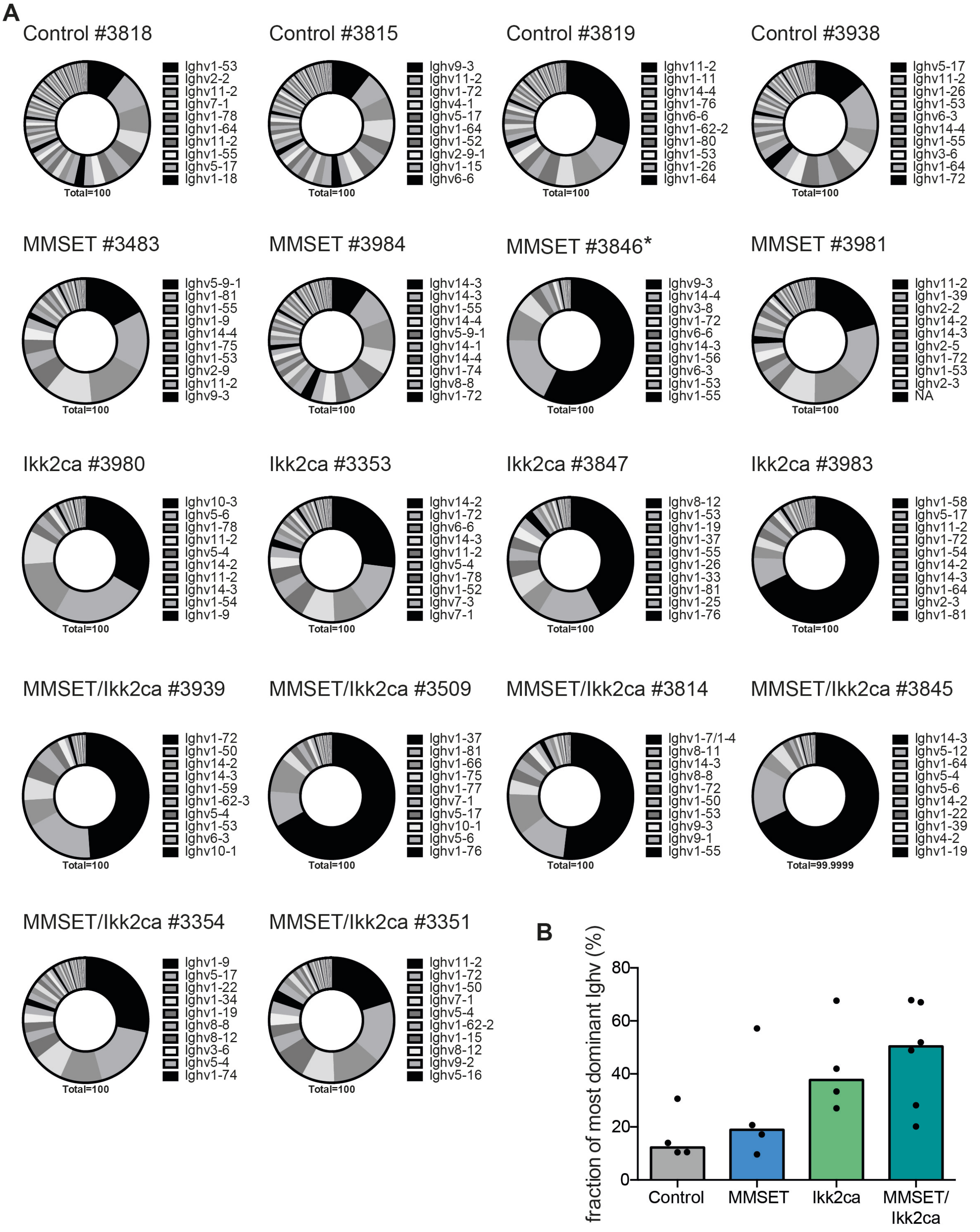

A
Control #3818
Ighv1-53
Ighv2-2
Ighv11-2
Ighv7-1
Ighv1-78
Ighv1-64
Ighv11-2
Ighv1-55
Ighv5-17
Ighv1-18
Total=100
Control #3815
Ighv9-3
Ighv11-2
Ighv1-72
Ighv4-1
Ighv5-17
Ighv1-64
Ighv1-52
Ighv2-9-1
Ighv1-15
Ighv6-6
Total=100
Control #3819
Ighv11-2
Ighv1-11
Ighv14-4
Ighv1-76
Ighv6-6
Ighv1-62-2
Ighv1-80
Ighv1-53
Ighv1-26
Ighv1-64
Total=100
Control #3938
Ighv5-17
Ighv11-2
Ighv1-26
Ighv1-53
Ighv6-3
Ighv14-4
Ighv1-55
Ighv3-6
Ighv1-64
Ighv1-72
Total=100
MMSET #3483
Ighv5-9-1
Ighv1-81
Ighv1-55
Ighv1-9
Ighv14-4
Ighv1-75
Ighv1-53
Ighv2-9
Ighv11-2
Ighv9-3
Total=100
MMSET #3984
Ighv14-3
Ighv14-3
Ighv1-55
Ighv14-4
Ighv5-9-1
Ighv14-1
Ighv14-4
Ighv1-74
Ighv8-8
Ighv1-72
Total=100
MMSET #3846*
Ighv9-3
Ighv14-4
Ighv3-8
Ighv1-72
Ighv6-6
Ighv14-3
Ighv1-56
Ighv6-3
Ighv1-53
Ighv1-55
Total=100
MMSET #3981
Ighv11-2
Ighv1-39
Ighv2-2
Ighv14-2
Ighv14-3
Ighv2-5
Ighv1-72
Ighv1-53
Ighv2-3
NA
Total=100
Ikk2ca #3980
Ighv10-3
Ighv5-6
Ighv1-78
Ighv11-2
Ighv5-4
Ighv14-2
Ighv11-2
Ighv14-3
Ighv1-54
Ighv1-9
Total=100
Ikk2ca #3353
Ighv14-2
Ighv1-72
Ighv6-6
Ighv14-3
Ighv11-2
Ighv5-4
Ighv1-78
Ighv1-52
Ighv7-3
Ighv7-1
Total=100
Ikk2ca #3847
Ighv8-12
Ighv1-53
Ighv1-19
Ighv1-37
Ighv1-55
Ighv1-26
Ighv1-33
Ighv1-81
Ighv1-25
Ighv1-76
Total=100
Ikk2ca #3983
Ighv1-58
Ighv5-17
Ighv11-2
Ighv1-72
Ighv1-54
Ighv14-2
Ighv14-3
Ighv1-64
Ighv2-3
Ighv1-81
Total=100
MMSET/Ikk2ca #3939
Ighv1-72
Ighv1-50
Ighv14-2
Ighv14-3
Ighv1-59
Ighv1-62-3
Ighv5-4
Ighv1-53
Ighv6-3
Ighv10-1
Total=100
MMSET/Ikk2ca #3509
Ighv1-37
Ighv1-81
Ighv1-66
Ighv1-75
Ighv1-77
Ighv7-1
Ighv5-17
Ighv10-1
Ighv5-6
Ighv1-76
Total=100
MMSET/Ikk2ca #3814
Ighv1-7/1-4
Ighv8-11
Ighv14-3
Ighv8-8
Ighv1-72
Ighv1-50
Ighv1-53
Ighv9-3
Ighv9-1
Ighv1-55
Total=100
MMSET/Ikk2ca #3845
Ighv14-3
Ighv5-12
Ighv1-64
Ighv5-4
Ighv5-6
Ighv14-2
Ighv1-22
Ighv1-39
Ighv4-2
Ighv1-19
Total=99.9999
MMSET/Ikk2ca #3354
Ighv1-9
Ighv5-17
Ighv1-22
Ighv1-34
Ighv1-19
Ighv8-8
Ighv8-12
Ighv3-6
Ighv5-4
Ighv1-74
Total=100
MMSET/Ikk2ca #3351
Ighv11-2
Ighv1-72
Ighv1-50
Ighv7-1
Ighv5-4
Ighv1-62-2
Ighv1-15
Ighv8-12
Ighv9-2
Ighv5-16
Total=100
B
fraction of most dominant Ighv (%)
80
60
40
20
0
Control
MMSET
Ikk2ca
MMSET/Ikk2ca

#### Figure S6

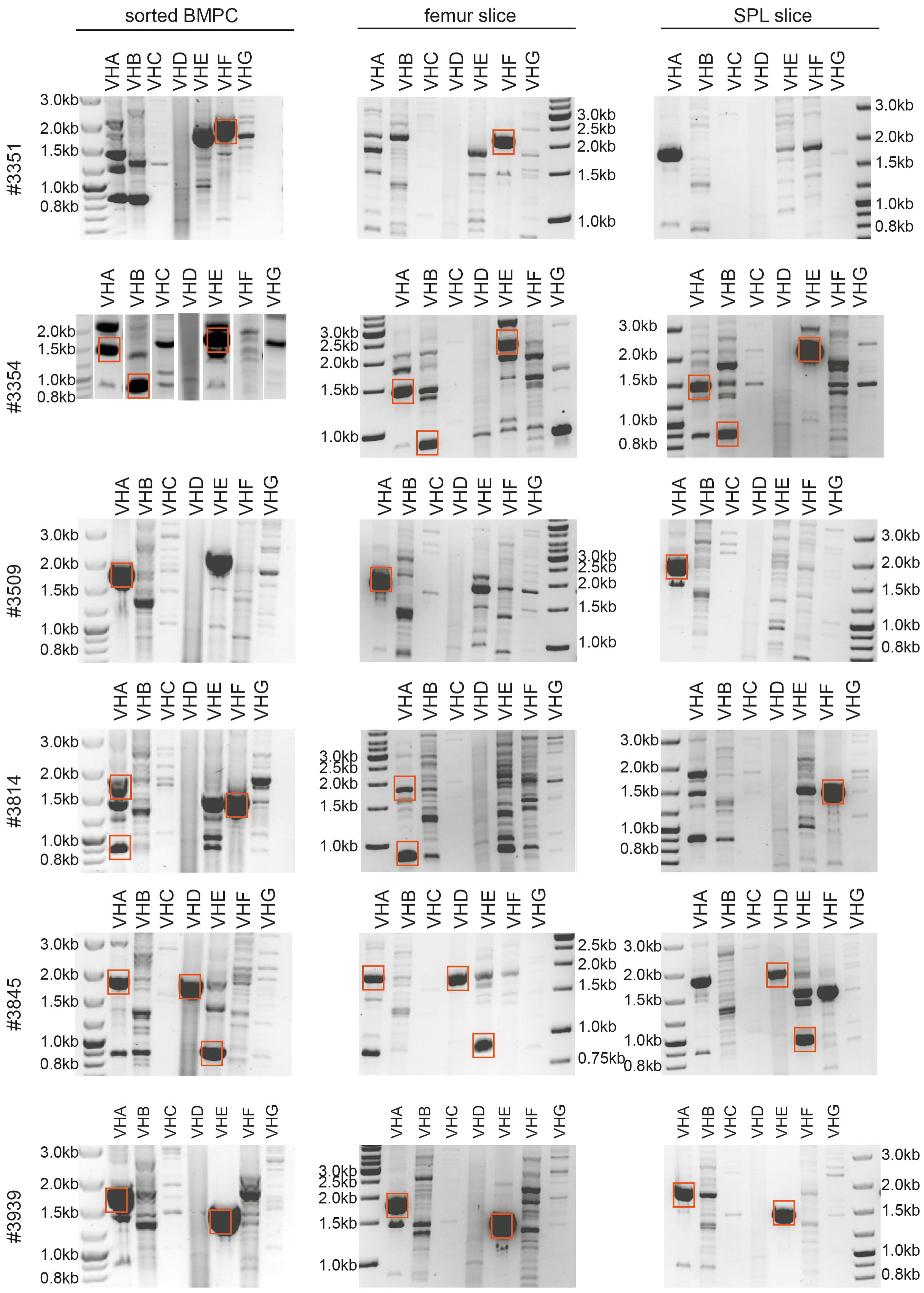
sorted BMPC
femur slice
SPL slice
#3351
#3354
#3509
#3814
#3845
#3939
VHA
VHB
VHC
VHD
VHE
VHF
VHG
3.0kb
2.5kb
2.0kb
1.5kb
1.0kb
0.8kb
0.75kb

#### Figure S7

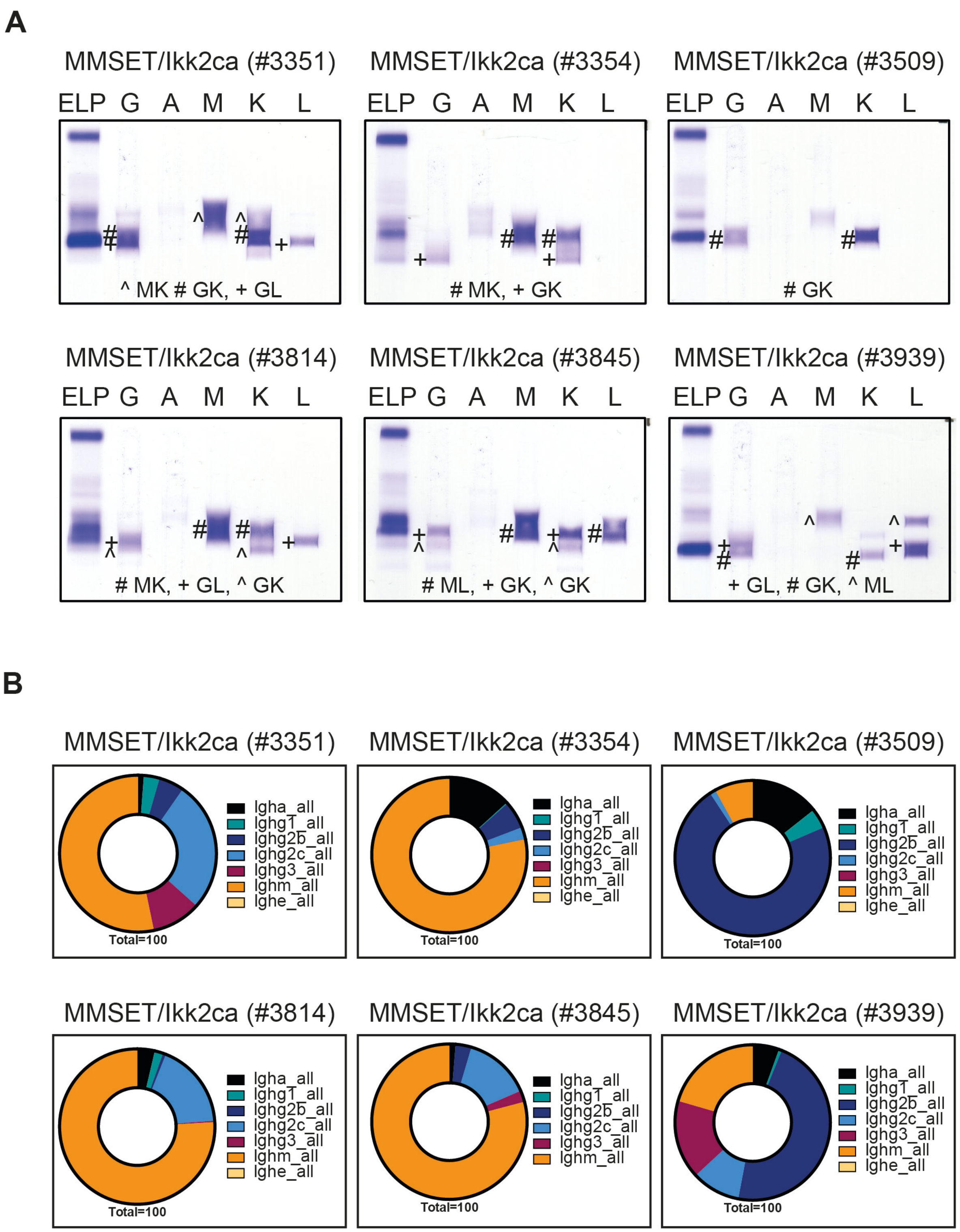
A
MMSET/Ikk2ca (#3351)
ELP G A M K L
^ MK # GK, + GL
MMSET/Ikk2ca (#3354)
ELP G A M K L
# MK, + GK
MMSET/Ikk2ca (#3509)
ELP G A M K L
# GK
MMSET/Ikk2ca (#3814)
ELP G A M K L
# MK, + GL, ^ GK
MMSET/Ikk2ca (#3845)
ELP G A M K L
# ML, + GK, ^ GK
MMSET/Ikk2ca (#3939)
ELP G A M K L
+ GL, # GK, ^ ML
B
MMSET/Ikk2ca (#3351)
Igha_all
Ighg1_all
Ighg2b_all
Ighg2c_all
Ighg3_all
Ighm_all
Ighe_all
Total=100
MMSET/Ikk2ca (#3354)
Igha_all
Ighg1_all
Ighg2b_all
Ighg2c_all
Ighg3_all
Ighm_all
Ighe_all
Total=100
MMSET/Ikk2ca (#3509)
Igha_all
Ighg1_all
Ighg2b_all
Ighg2c_all
Ighg3_all
Ighm_all
Ighe_all
Total=100
MMSET/Ikk2ca (#3814)
Igha_all
Ighg1_all
Ighg2b_all
Ighg2c_all
Ighg3_all
Ighm_all
Ighe_all
Total=100
MMSET/Ikk2ca (#3845)
Igha_all
Ighg1_all
Ighg2b_all
Ighg2c_all
Ighg3_all
Ighm_all
Total=100
MMSET/Ikk2ca (#3939)
Igha_all
Ighg1_all
Ighg2b_all
Ighg2c_all
Ighg3_all
Ighm_all
Ighe_all
Total=100

**Figure S8**

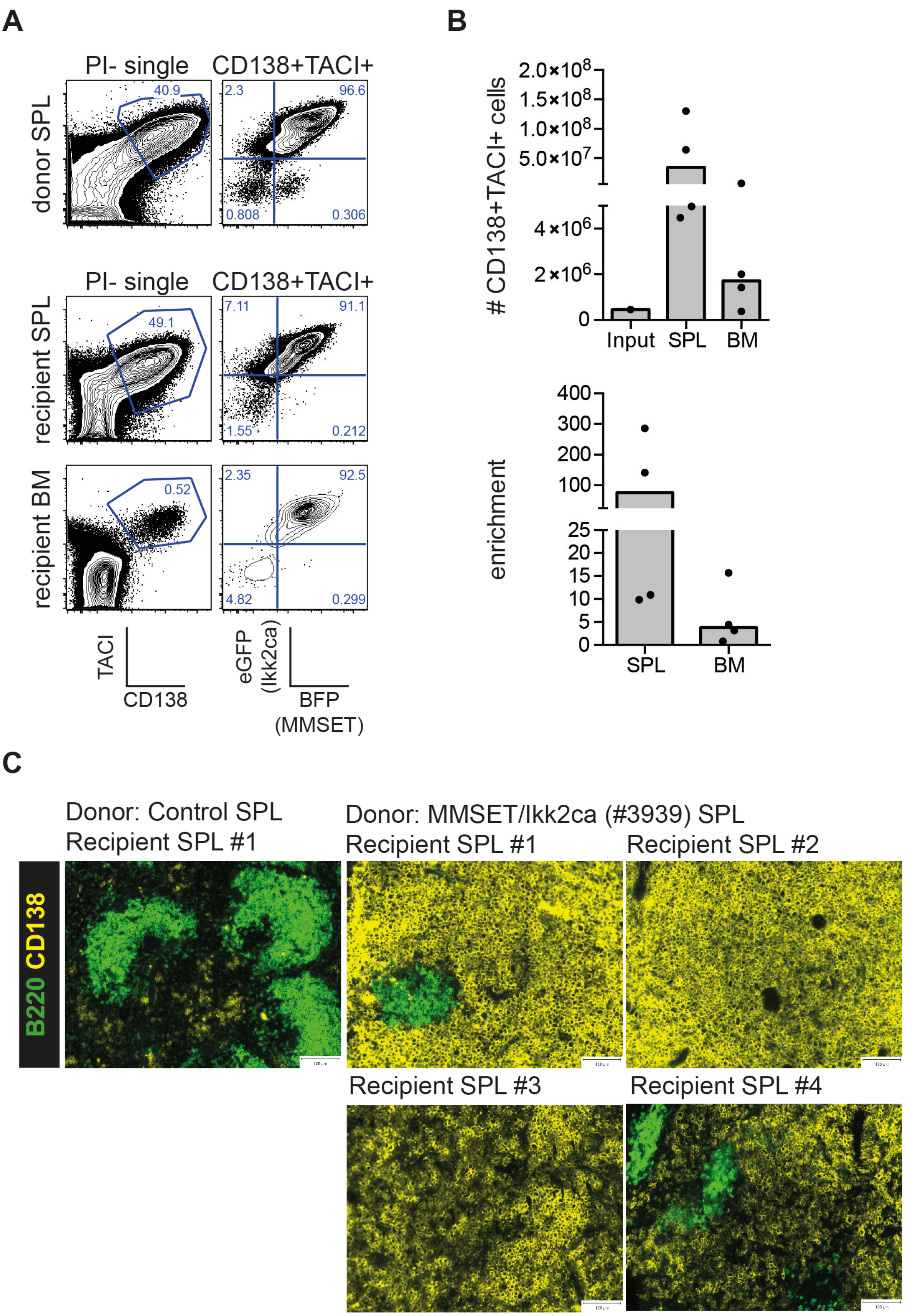
A
PI- single
CD138+TACI+
donor SPL
40.9
2.3
96.6
0.808
0.306
recipient SPL
49.1
7.11
91.1
1.55
0.212
recipient BM
0.52
2.35
92.5
4.82
0.299
TACI
CD138
eGFP (Ikk2ca)
BFP (MMSET)
B
### CD138+TACI+ cells
2.0×10^8
1.5×10^8
1.0×10^8
5.0×10^7
4×10^6
2×10^6
0
Input
SPL
BM
enrichment
400
300
200
100
25
20
15
10
5
0
SPL
BM
C
Donor: Control SPL
Recipient SPL #1
Donor: MMSET/Ikk2ca (#3939) SPL
Recipient SPL #1
Recipient SPL #2
Recipient SPL #3
Recipient SPL #4
B220 CD138

#### Figure S9

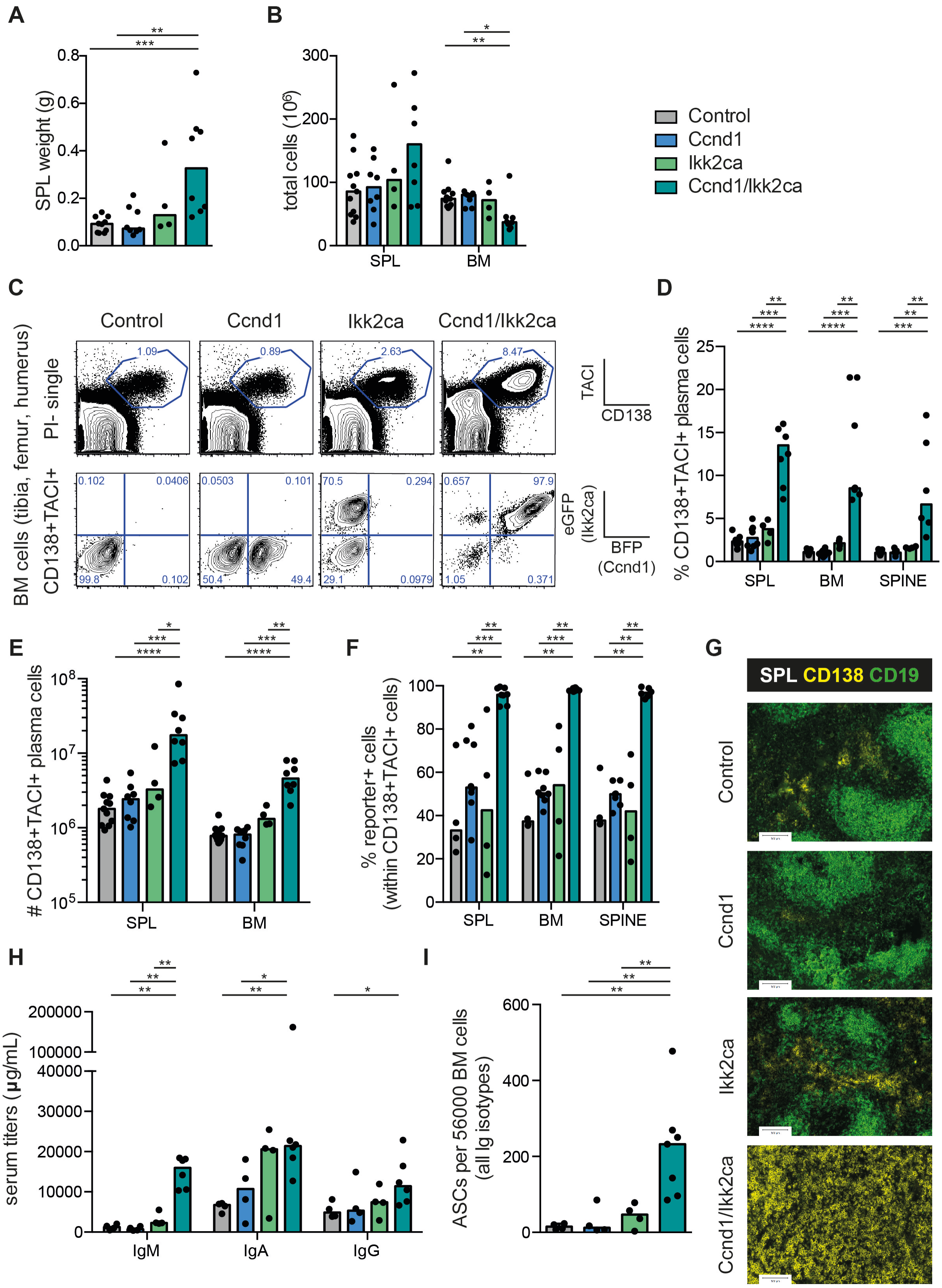
A
SPL weight (g)
B
total cells ($10^6$)
SPL
BM
Control
Ccnd1
Ikk2ca
Ccnd1/Ikk2ca
C
Control
Ccnd1
Ikk2ca
Ccnd1/Ikk2ca
BM cells (tibia, femur, humerus)
PI- single
CD138+TACI+
TACI
CD138
eGFP (Ikk2ca)
BFP (Ccnd1)
D
% CD138+TACI+ plasma cells
SPL
BM
SPINE
E
### CD138+TACI+ plasma cells
F
% reporter+ cells (within CD138+TACI+ cells)
G
SPL CD138 CD19
H
serum titers (µg/mL)
IgM
IgA
IgG
I
ASCs per 56000 BM cells (all Ig isotypes)

### Figure S10

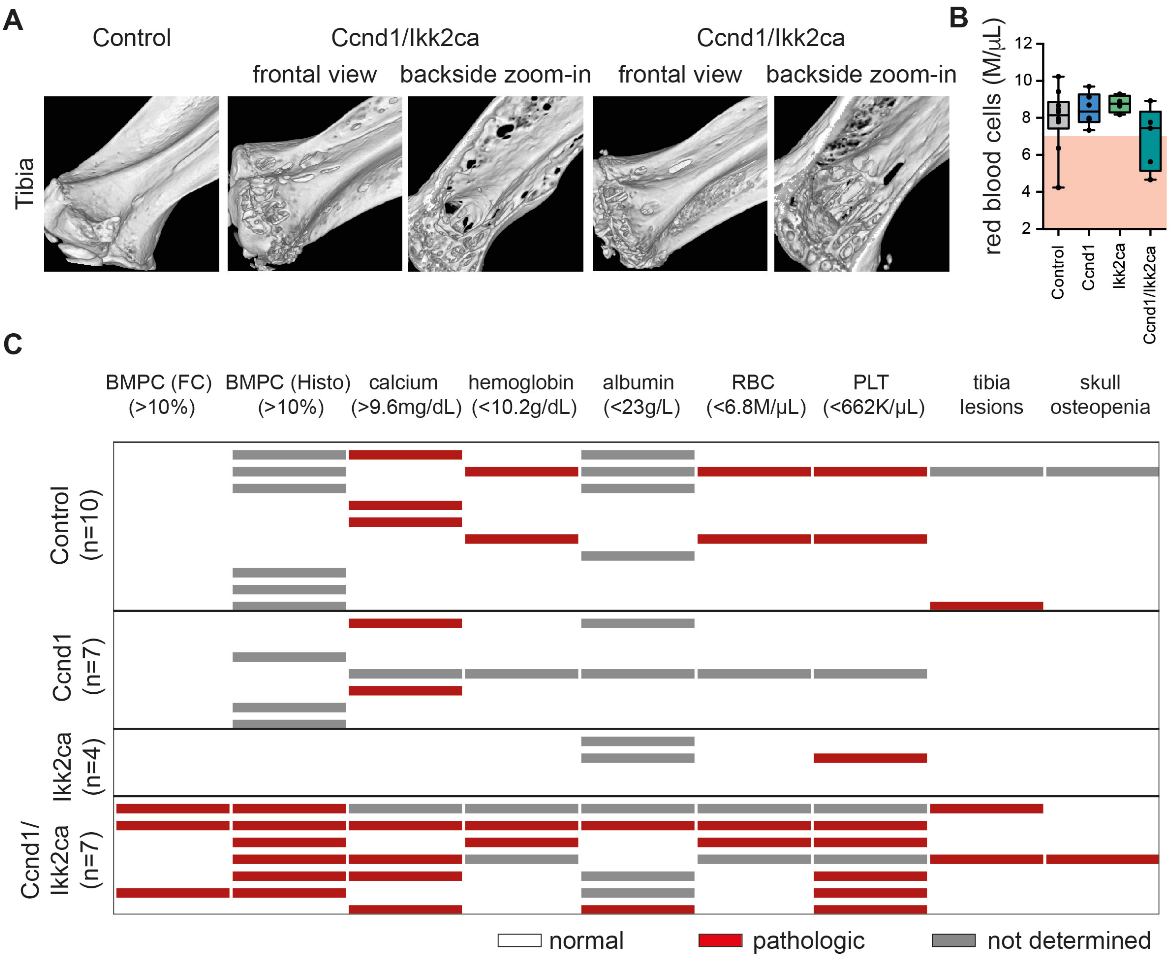

A
Control
Ccnd1/Ikk2ca
frontal view
backside zoom-in
Ccnd1/Ikk2ca
frontal view
backside zoom-in
Tibia
B
red blood cells (M/μL)
12
10
8
6
4
2
Control
Ccnd1
Ikk2ca
Ccnd1/Ikk2ca
C
BMPC (FC) (>10%)
BMPC (Histo) (>10%)
calcium (>9.6mg/dL)
hemoglobin (<10.2g/dL)
albumin (<23g/L)
RBC (<6.8M/μL)
PLT (<662K/μL)
tibia lesions
skull osteopenia
Control (n=10)
Ccnd1 (n=7)
Ikk2ca (n=4)
Ccnd1/ Ikk2ca (n=7)
normal
pathologic
not determined

### Figure S11

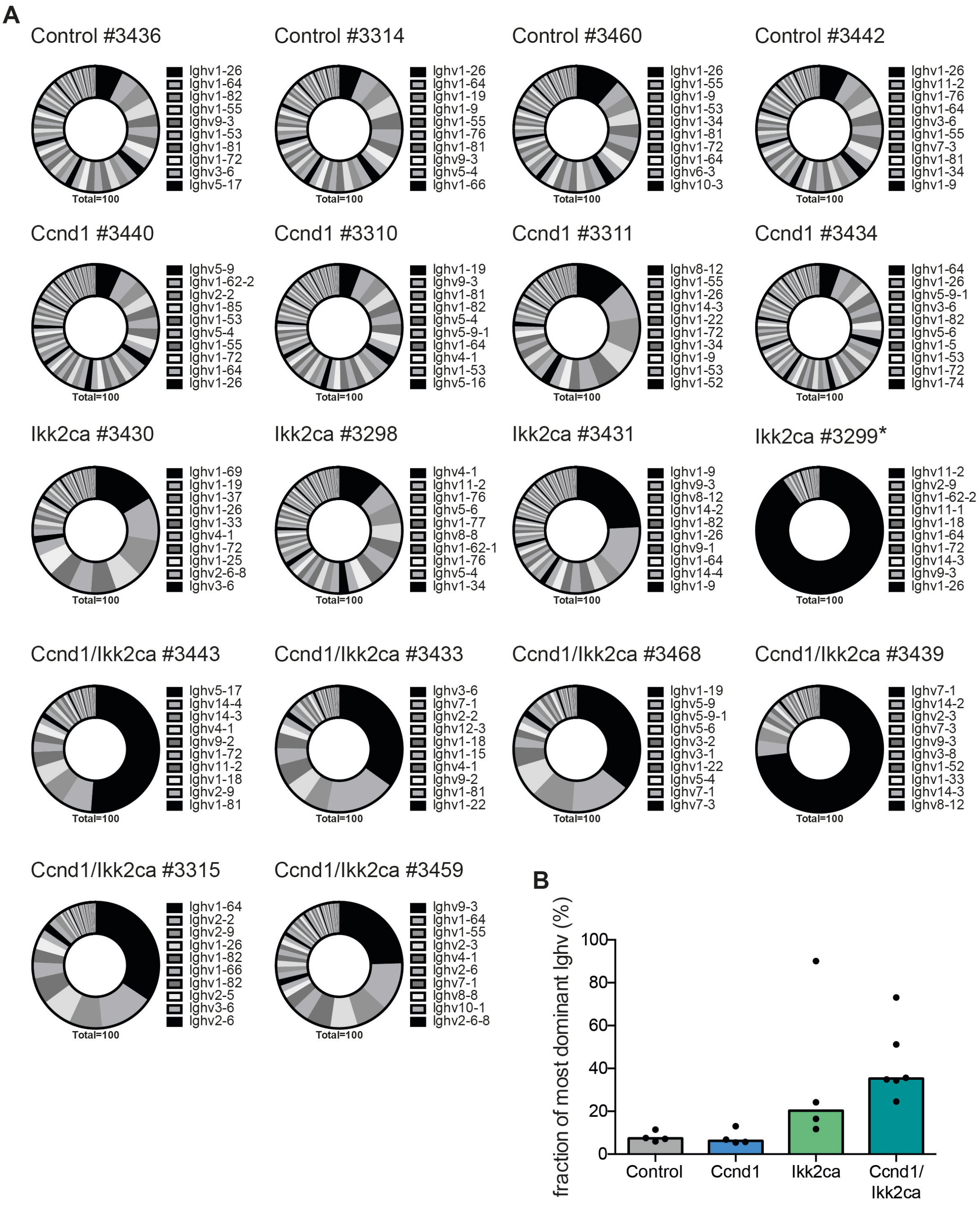
A
Control #3436
Ighv1-26
Ighv1-64
Ighv1-82
Ighv1-55
Ighv9-3
Ighv1-53
Ighv1-81
Ighv1-72
Ighv3-6
Ighv5-17
Total=100
Control #3314
Ighv1-26
Ighv1-64
Ighv1-19
Ighv1-9
Ighv1-55
Ighv1-76
Ighv1-81
Ighv9-3
Ighv5-4
Ighv1-66
Total=100
Control #3460
Ighv1-26
Ighv1-55
Ighv1-9
Ighv1-53
Ighv1-34
Ighv1-81
Ighv1-72
Ighv1-64
Ighv6-3
Ighv10-3
Total=100
Control #3442
Ighv1-26
Ighv11-2
Ighv1-76
Ighv1-64
Ighv3-6
Ighv1-55
Ighv7-3
Ighv1-81
Ighv1-34
Ighv1-9
Total=100
Ccnd1 #3440
Ighv5-9
Ighv1-62-2
Ighv2-2
Ighv1-85
Ighv1-53
Ighv5-4
Ighv1-55
Ighv1-72
Ighv1-64
Ighv1-26
Total=100
Ccnd1 #3310
Ighv1-19
Ighv9-3
Ighv1-81
Ighv1-82
Ighv5-4
Ighv5-9-1
Ighv1-64
Ighv4-1
Ighv1-53
Ighv5-16
Total=100
Ccnd1 #3311
Ighv8-12
Ighv1-55
Ighv1-26
Ighv14-3
Ighv1-22
Ighv1-72
Ighv1-34
Ighv1-9
Ighv1-53
Ighv1-52
Total=100
Ccnd1 #3434
Ighv1-64
Ighv1-26
Ighv5-9-1
Ighv3-6
Ighv1-82
Ighv5-6
Ighv1-5
Ighv1-53
Ighv1-72
Ighv1-74
Total=100
Ikk2ca #3430
Ighv1-69
Ighv1-19
Ighv1-37
Ighv1-26
Ighv1-33
Ighv4-1
Ighv1-72
Ighv1-25
Ighv2-6-8
Ighv3-6
Total=100
Ikk2ca #3298
Ighv4-1
Ighv11-2
Ighv1-76
Ighv5-6
Ighv1-77
Ighv8-8
Ighv1-62-1
Ighv1-76
Ighv5-4
Ighv1-34
Total=100
Ikk2ca #3431
Ighv1-9
Ighv9-3
Ighv8-12
Ighv14-2
Ighv1-82
Ighv1-26
Ighv9-1
Ighv1-64
Ighv14-4
Ighv1-9
Total=100
Ikk2ca #3299*
Ighv11-2
Ighv2-9
Ighv1-62-2
Ighv11-1
Ighv1-18
Ighv1-64
Ighv1-72
Ighv14-3
Ighv9-3
Ighv1-26
Total=100
Ccnd1/Ikk2ca #3443
Ighv5-17
Ighv14-4
Ighv14-3
Ighv4-1
Ighv9-2
Ighv1-72
Ighv11-2
Ighv1-18
Ighv2-9
Ighv1-81
Total=100
Ccnd1/Ikk2ca #3433
Ighv3-6
Ighv7-1
Ighv2-2
Ighv12-3
Ighv1-18
Ighv1-15
Ighv4-1
Ighv9-2
Ighv1-81
Ighv1-22
Total=100
Ccnd1/Ikk2ca #3468
Ighv1-19
Ighv5-9
Ighv5-9-1
Ighv5-6
Ighv3-2
Ighv3-1
Ighv1-22
Ighv5-4
Ighv7-1
Ighv7-3
Total=100
Ccnd1/Ikk2ca #3439
Ighv7-1
Ighv14-2
Ighv2-3
Ighv7-3
Ighv9-3
Ighv3-8
Ighv1-52
Ighv1-33
Ighv14-3
Ighv8-12
Total=100
Ccnd1/Ikk2ca #3315
Ighv1-64
Ighv2-2
Ighv2-9
Ighv1-26
Ighv1-82
Ighv1-66
Ighv1-82
Ighv2-5
Ighv3-6
Ighv2-6
Total=100
Ccnd1/Ikk2ca #3459
Ighv9-3
Ighv1-64
Ighv1-55
Ighv2-3
Ighv4-1
Ighv2-6
Ighv7-1
Ighv8-8
Ighv10-1
Ighv2-6-8
Total=100
B
fraction of most dominant Ighv (%)
100
80
60
40
20
0
Control
Ccnd1
Ikk2ca
Ccnd1/
Ikk2ca

**Figure S12**

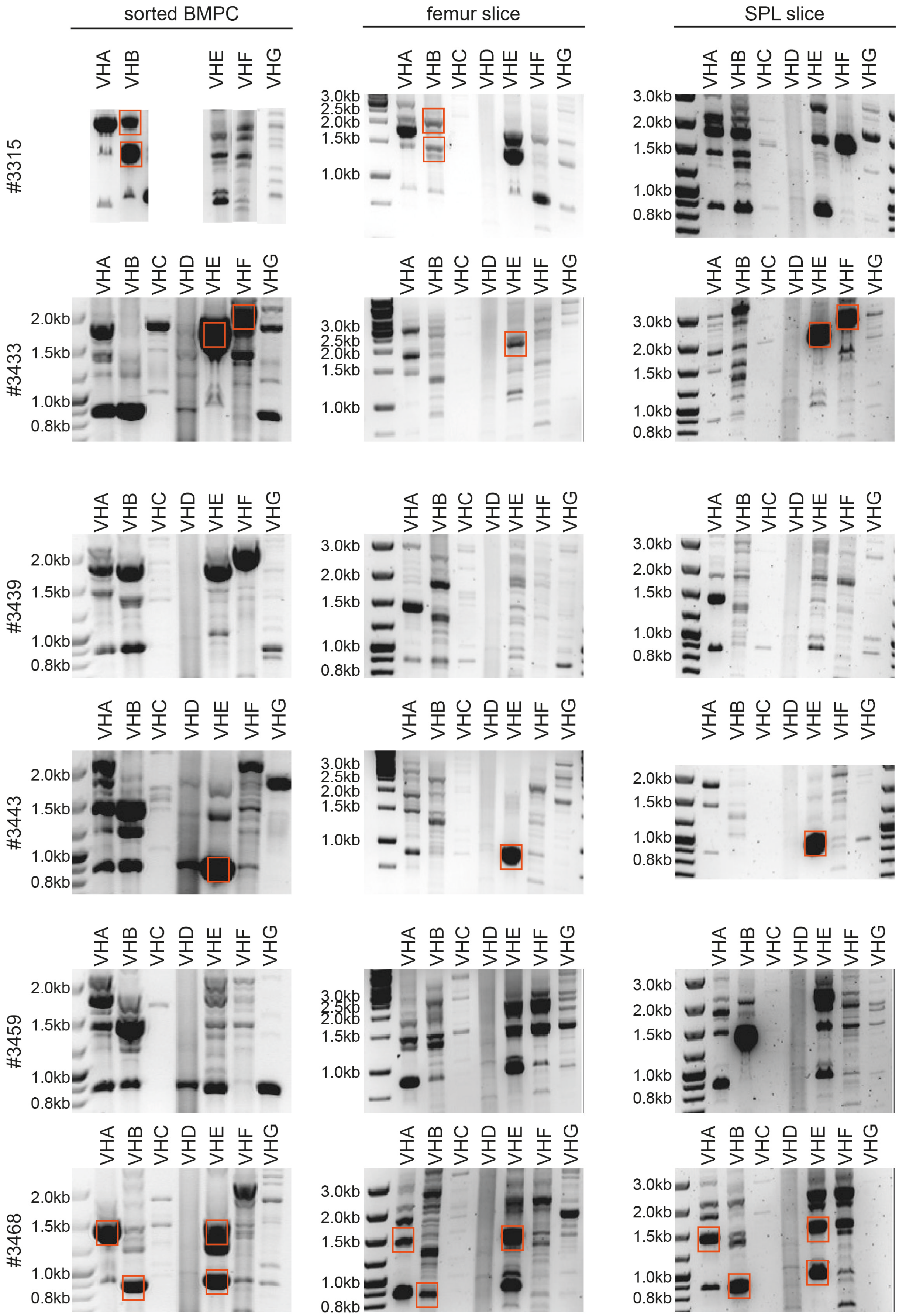
sorted BMPC
femur slice
SPL slice
#3315
#3433
#3439
#3443
#3459
#3468
VHA
VHB
VHC
VHD
VHE
VHF
VHG
3.0kb
2.5kb
2.0kb
1.5kb
1.0kb
0.8kb

#### Figure S13

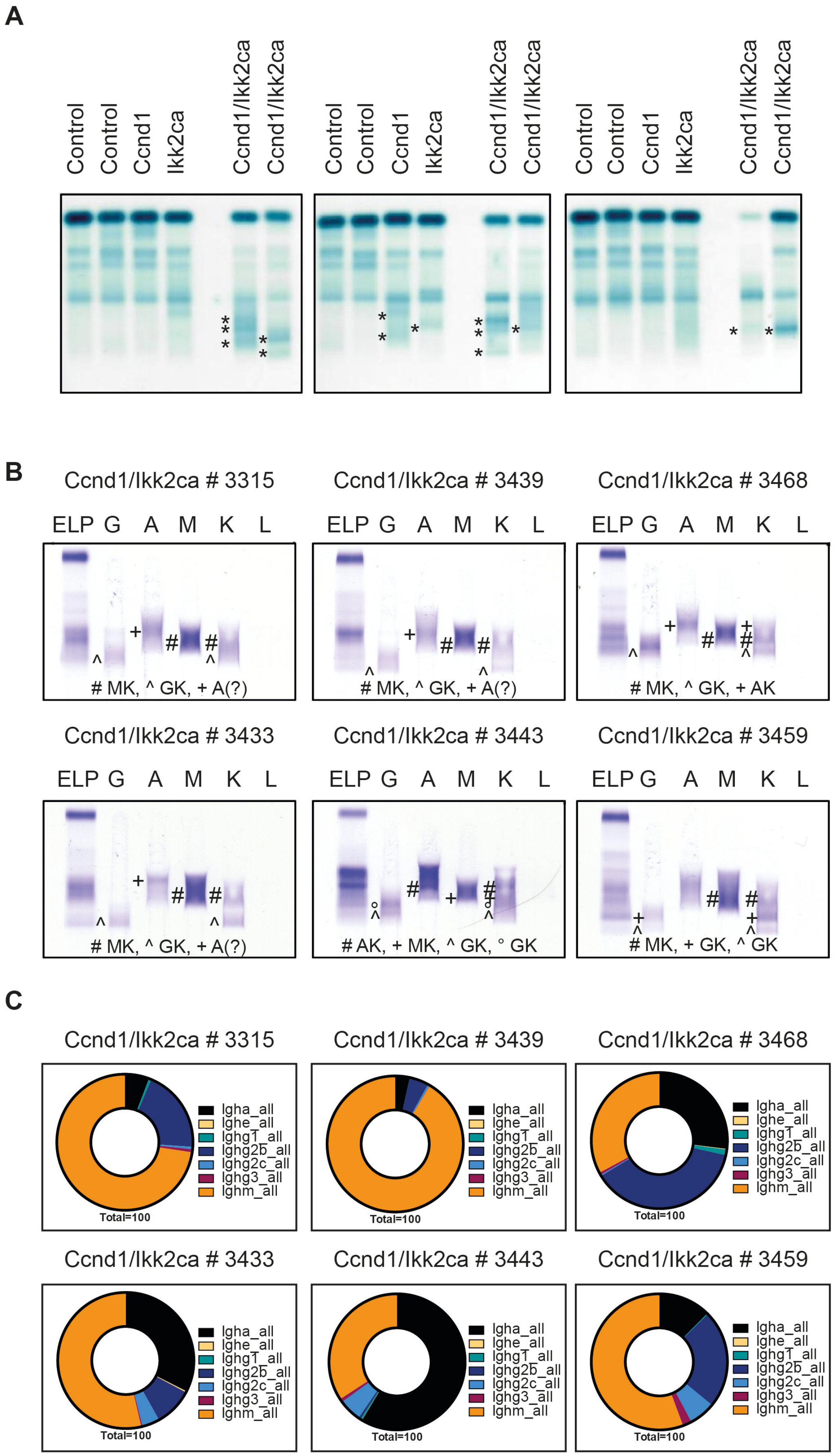
A
Control
Control
Ccnd1
Ikk2ca
Ccnd1/Ikk2ca
Ccnd1/Ikk2ca
B
Ccnd1/Ikk2ca # 3315
Ccnd1/Ikk2ca # 3439
Ccnd1/Ikk2ca # 3468
ELP G A M K L
# MK, ^ GK, + A(?)
# MK, ^ GK, + AK
Ccnd1/Ikk2ca # 3433
Ccnd1/Ikk2ca # 3443
Ccnd1/Ikk2ca # 3459
# AK, + MK, ^ GK, ° GK
# MK, + GK, ^ GK
C
Igha_all
Ighe_all
Ighg1_all
Ighg2b_all
Ighg2c_all
Ighg3_all
Ighm_all
Total=100

**Figure S14**

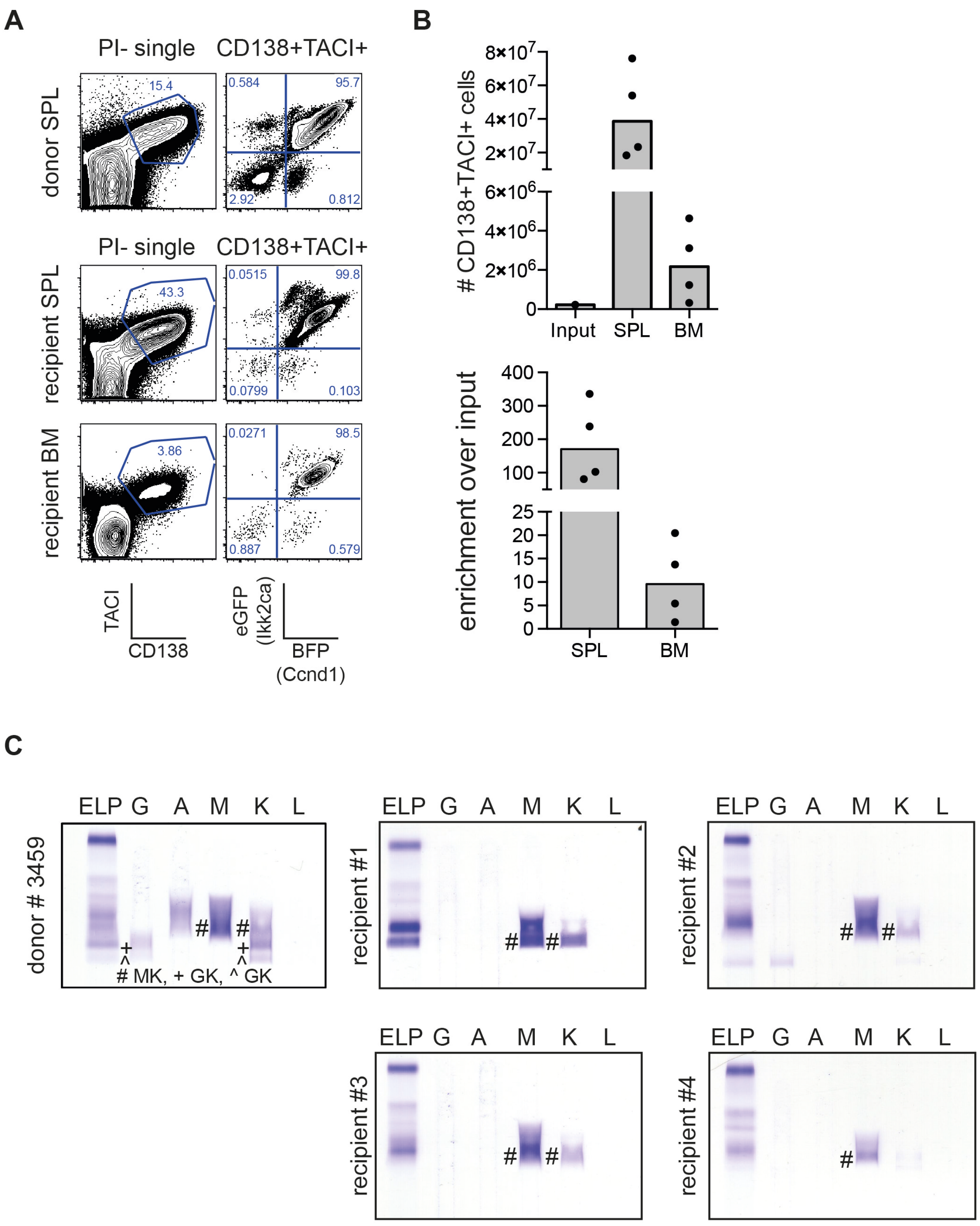

A
PI- single
CD138+TACI+
donor SPL
recipient SPL
recipient BM
15.4
0.584
95.7
2.92
0.812
43.3
0.0515
99.8
0.0799
0.103
3.86
0.0271
98.5
0.887
0.579
TACI
CD138
eGFP (Ikk2ca)
BFP (Ccnd1)
B
### CD138+TACI+ cells
Input
SPL
BM
enrichment over input
C
ELP G A M K L
donor # 3459
# MK, + GK, ^ GK
recipient #1
recipient #2
recipient #3
recipient #4

### Figure S15

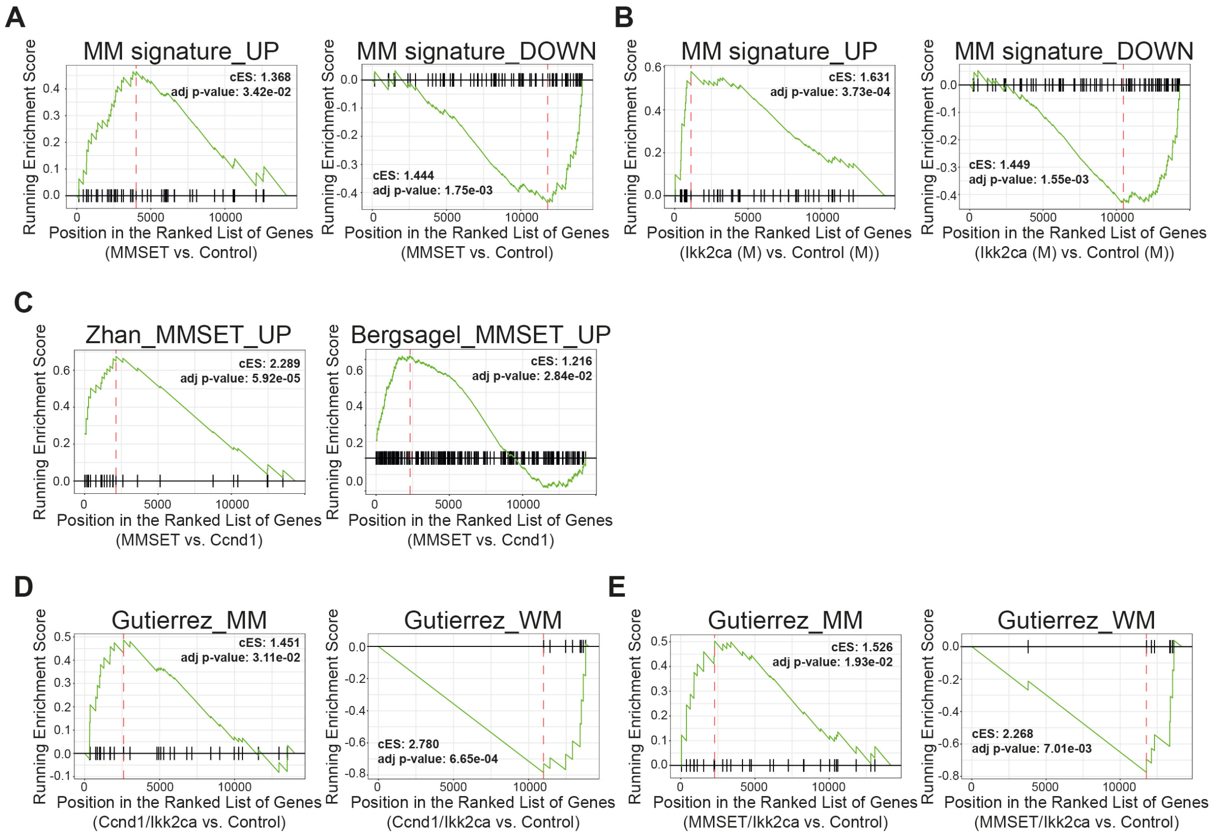
A
MM signature_UP
cES: 1.368
adj p-value: 3.42e-02
Position in the Ranked List of Genes
(MMSET vs. Control)
MM signature_DOWN
cES: 1.444
adj p-value: 1.75e-03
Position in the Ranked List of Genes
(MMSET vs. Control)
B
MM signature_UP
cES: 1.631
adj p-value: 3.73e-04
Position in the Ranked List of Genes
(Ikk2ca (M) vs. Control (M))
MM signature_DOWN
cES: 1.449
adj p-value: 1.55e-03
Position in the Ranked List of Genes
(Ikk2ca (M) vs. Control (M))
C
Zhan_MMSET_UP
cES: 2.289
adj p-value: 5.92e-05
Position in the Ranked List of Genes
(MMSET vs. Ccnd1)
Bergsagel_MMSET_UP
cES: 1.216
adj p-value: 2.84e-02
Position in the Ranked List of Genes
(MMSET vs. Ccnd1)
D
Gutierrez_MM
cES: 1.451
adj p-value: 3.11e-02
Position in the Ranked List of Genes
(Ccnd1/Ikk2ca vs. Control)
Gutierrez_WM
cES: 2.780
adj p-value: 6.65e-04
Position in the Ranked List of Genes
(Ccnd1/Ikk2ca vs. Control)
E
Gutierrez_MM
cES: 1.526
adj p-value: 1.93e-02
Position in the Ranked List of Genes
(MMSET/Ikk2ca vs. Control)
Gutierrez_WM
cES: 2.268
adj p-value: 7.01e-03
Position in the Ranked List of Genes
(MMSET/Ikk2ca vs. Control)
Running Enrichment Score

#### Supplementary Table Legends

**Table S1: Histopathologic evaluation of femur and spleen sections from MMSET cohort mice.** Sections from spleens and femurs of aged control, single mutant (MMSET, Ikk2ca) and double mutant (MMSET/Ikk2ca) mice were stained with HE and evaluated in a blinded fashion without knowledge of the underlying genotype. ND, denotes not determined.

**Table S2: Overview of dominant VDJ rearrangements of MMSET/Ikk2ca plasma cells determined by VDJ-PCR.** Abbreviations: P/NP = productive/non-productive rearrangement; tissue = SPL (spleen section), F (femur section), BMPC (sorted (reporter+) CD138+TACI+ long bone-derived BM plasma cells); #SHM = amount of somatic hypermutation within VDJ rearrangement.

**Table S3: Histopathologic evaluation of femur and spleen sections from Ccnd1 cohort mice.** Sections from spleens and femurs of aged control, single mutant (Ccnd1, Ikk2ca) and double mutant (Ccnd1/Ikk2ca) mice were stained with HE and evaluated in a blinded fashion without knowledge of the underlying genotype.

**Table S4: Overview of dominant VDJ rearrangements of Ccnd1/Ikk2ca plasma cells determined by VDJ-PCR.** Abbreviations: P/NP = productive/non-productive rearrangement; tissue = SPL (spleen section), F (femur section), BMPC (sorted (reporter+) CD138+TACI+ long bone-derived BM plasma cells); #SHM = amount of somatic hypermutation within VDJ rearrangement.

**Table S5: List of Antibodies**

**Table S6: Overview of gene sets tested in the GSEA**

**Table S1**

| mouse ID | genotype | Femur | Spleen |
|---|---|---|---|
| 3815 | control | ND | enrichment of plasma cells |
| 3818 | control | normal | plasma cell hyperplasia in marginal zone (no tumor) |
| 3819 | control | normal | plasma cell hyperplasia in marginal zone (no tumor) |
| 3938 | control | normal | plasma cell hyperplasia in marginal zone (no tumor) |
| 5230 | control | normal | ND |
| 4978 | control | normal | ND |
| 4977 | control | normal | ND |
| 3483 | MMSET | normal | plasma cell hyperplasia in marginal zone (no tumor) |
| 3846 | MMSET | diffuse myeloma infiltrate (99 %) | tumor-forming plasma cell infiltrates, destruction of architecture |
| 3981 | MMSET | normal | plasma cell hyperplasia in marginal zone (no tumor) |
| 3984 | MMSET | normal | plasma cell hyperplasia in marginal zone (no tumor) |
| 5250 | MMSET | normal | ND |
| 5253 | MMSET | normal | ND |
| 4979 | MMSET | normal | ND |
| 3353 | Ikk2ca | diffuse myeloma infiltrate (70 %) | tumor-forming plasma cell infiltrates, destruction of architecture |
| 3847 | Ikk2ca | focal myeloma infiltrate (approx. 5 %) | tumor-forming plasma cell infiltrates, destruction of architecture |
| 3980 | Ikk2ca | normal | nodular plasma cell infiltrates |
| 3983 | Ikk2ca | normal | plasma cell hyperplasia in marginal zone (no tumor) |
| 5250 | Ikk2ca | normal | ND |
| 5643 | Ikk2ca | normal | ND |
| 5161 | Ikk2ca | normal | ND |
| 5261 | Ikk2ca | normal | ND |

| | | | |
|---|---|---|---|
| 3351 | MMSET/Ikk2ca | diffuse myeloma infiltrate (90 %) | tumor-forming plasma cell infiltrates, destruction of architecture |
| 3354 | MMSET/Ikk2ca | diffuse myeloma infiltrate (approx. 80 %) | tumor-forming plasma cell infiltrates, destruction of architecture |
| 3509 | MMSET/Ikk2ca | diffuse myeloma infiltrate (90 %) | tumor-forming plasma cell infiltrates, destruction of architecture |
| 3814 | MMSET/Ikk2ca | diffuse myeloma infiltrate (99 %) | tumor-forming plasma cell infiltrates, destruction of architecture |
| 3845 | MMSET/Ikk2ca | diffuse myeloma infiltrate (90 %) | tumor-forming plasma cell infiltrates, destruction of architecture |
| 3939 | MMSET/Ikk2ca | diffuse myeloma infiltrate (99 %) | tumor-forming plasma cell infiltrates, destruction of architecture |
| 5409 | MMSET/Ikk2ca | diffuse myeloma infiltrate (40 %)*** | ND |
| 5160 | MMSET/Ikk2ca | diffuse myeloma infiltrate (80 %) | ND |
| 5258 | MMSET/Ikk2ca | normal | ND |
| 5408 | MMSET/Ikk2ca | focal myeloma infiltrate (10%)* | ND |
| 4997 | MMSET/Ikk2ca | focal myeloma infiltrate (20%)** | ND |
| 5434 | MMSET/Ikk2ca | diffuse myeloma infiltrate (50 %)**** | ND |
| 4995 | MMSET/Ikk2ca | diffuse myeloma infiltrate (90 %) | ND |
| 5075 | MMSET/Ikk2ca | diffuse myeloma infiltrate (80 %) | ND |
| 5259 | MMSET/Ikk2ca | diffuse myeloma infiltrate (90 %) | ND |

NOTE: Most MMSET/Ikk2ca mice that did not show 60% myeloma infiltration in the histologic femur biopsy still presented with MM-associated clinical features reaching a myeloma-defining event (IMWG criteria; Rajkumar, et al., 2014).

* 5408: high calcium (9.93mg/dL), low albumin (22.36g/L), low RBCs (6.74M/µL), low platelets (49K/µL), tibia lesions

** 4997: increased calcium (9.73mg/dL), low hemoglobin (6.80g/dL), low albumin (20.35g/L), low RBCs (4.05M/µL), low platelets (93K/µL)

*** 5409: increased calcium (9.67mg/dL), low hemoglobin (7.9g/dL), low albumin (15.22g/L), low RBCs (5.56M/µL), low platelets (384K/µL)

**** 5434: low hemoglobin (8.00g/dL), low albumin (13.25g/L), low RBCs (4.87M/µL), low platelets (231K/µL), tibia lesions, skull osteopenia

**Table S2**

| mouse ID | genotype | PCR fragment | VDJ rearrangement | CDR3 | P/ NP | tissue | # SHM |
|---|---|---|---|---|---|---|---|
| 3354 | MMSET/ Ikk2ca | VHE-JH1 | Ighv5-17*01; Ighd1-1*01; Ighj1*03 | GCAAGGCCCTTTACTACGATAGGAGCTACGTACTGGTACTTCGATGTC | P | SPL, F, BMPC | 0 |
| | | VHB-JH4 | Ighv2-6-5*01; Ighd2-1*01; Ighj4*01 | GCCAAATACTATGGTAACTACTATGCTATGGACTAC | P | SPL, F, BMPC | 5 to 8 |
| | | VHA-JH3 | Ighv1-37*01; Ighd2-1*01; Ighj3*01 | GCAAGATTTCTCAGTCTACTATGGTAACGGTGGGCCTGGTTTGCTTAC | NP | SPL, F, BMPC | 0 |
| 3814 | MMSET/ Ikk2ca | VHF-JH3 | Ighv6-6*01; Ighd2-1*01; Ighj3*01 | ACCAGGCCGGGCTATGGTAACCCCTGGTTTGCTTAC | P | SPL, BMPC | 1 to 3 |
| | | VHA-JH2 | Ighv1-4*01/Ighv1-7*01; Ighd1-1*01; Ighj2*01 | GCAACCCCTTATTACTACGGTAGTAGCTACAACTAC | P | SPL, F, BMPC | 5 to 8 |
| | | VHA-JH4 | Ighv1-81*01/Ighd1-2*01/Ighj4*01 | GCAAGAGGTATTACTACGGCTACTACTATGCTATGGACTAC | NP | F, BMPC | 13 to 15 |
| 3845 | MMSET/ Ikk2ca | VHE-JH4 | Ighv5-12*01; Ighd2-4*01; Ighj4*01 | GCAAGACATGATTACGAGGGGCCTAACTATGCTATGGACTAC | P | SPL, F, BMPC | 0 to 1 |
| | | VHA-JH2 | Ighv1s135*01/Ighd2-14*01/Ighj2*01 | GCAAGAGAGGGGGAAAACTTGAGGTACGACGAGGGAGGGGTTGACTAC | P | F, BMPC | 1 to 2 |
| | | VHD-JH1 | Ighv4-2*01; N/A; Ighj2*01 | GCAAGACAGGGGTACTACTTTGACTAC | NP | SPL, F, BMPC | 0 to 2 |
| | | Ighv14-3-JH4 | Ighv14-3*02; Ighd2-14*01; Ighj4*01 | GCTAGATACTATAGGTACCTCTATGCTATGGACTAC | P | F | 0 |
| 3939 | MMSET/ Ikk2ca | VHE-JH3 | Ighv5-4*02; Ighd2-14*01; Ighj3*01 | GCAAGAGATCGGTTCCCAATAGGTACGACGTTTTGCTTAC | NP | SPL, F, BMPC | 0 to 1 |
| | | VHA-JH2 | Ighv1-72*01; Ighd3-2*02; Ighj2*01 | GCAAGAGGACAGCTCAGGCTTCACTTTGACTAC | P | SPL, F, BMPC | 0 |
| 3509 | MMSET/ Ikk2ca | VHA-JH2 | Ighv1-37*01; Ighd2-4*01; Ighj2*01 | GCAAGAGGGGGGAGCTATGATTACGACAGGATGTACTTTGACTAC | P | SPL, BMPC | 0 to 2 |
| 3351 | MMSET/ Ikk2ca | VHF-JH1 | Ighv11-2*02; Ighd2-1*01; Ighj1*01 | ATGAGATATGGTAACTACTGGTACTTCGATGTC | P | F, BMPC | 0 |
| | | VHA-JH1 | Ighv1-72*01; Ighd2-3*01; Ighj2*01 | GCTCGCTATGATGGTTACCTTTTTGACTAC | P | SPL | 0 |

**Table S3**

| mouse ID | genotype | Femur | Spleen |
|---|---|---|---|
| 3314 | control | normal | normal |
| 3436 | control | normal | normal |
| 3442 | control | normal | normal |
| 3460 | control | normal | normal |
| 3310 | Ccnd1 | normal | normal |
| 3311 | Ccnd1 | normal | normal |
| 3434 | Ccnd1 | normal | normal |
| 3440 | Ccnd1 | normal | normal |
| 3298 | Ikk2ca | normal | normal |
| 3299 | Ikk2ca | normal | normal |
| 3430 | Ikk2ca | normal | myeloma infiltrate |
| 3431 | Ikk2ca | normal | normal |
| 3315 | Ccnd1/Ikk2ca | myeloma infiltrate (80 %) | extensive myeloma infiltrate in red pulp |
| 3433 | Ccnd1/Ikk2ca | myeloma infiltrate (80 %),<br>signs of bone remodeling | extensive myeloma infiltrate<br>w/ destruction of white pulp |
| 3439 | Ccnd1/Ikk2ca | normal | myeloma infiltrate |
| 3443 | Ccnd1/Ikk2ca | diffuse myeloma infiltrate (80 %) | myeloma infiltrate |
| 3444 | Ccnd1/Ikk2ca | myeloma infiltrate (60 %) | myeloma infiltrate |
| 3459 | Ccnd1/Ikk2ca | myeloma infiltrate (70 %) | extensive myeloma infiltrate in red pulp |
| 3468 | Ccnd1/Ikk2ca | myeloma infiltrate (50 %) | extensive myeloma infiltrate in red pulp |

NOTE: Ccnd1/Ikk2ca mouse #3468 did not show ≥ 60 % myeloma infiltration in the histologic femur biopsy, but presented with high serum calcium (9.97mg/dL) and low platelet (560K/µL) levels reaching a myeloma-defining event (IMWG criteria; Rajkumar, et al., 2014).

**Table S4**

| mouse ID | genotype | PCR fragment | VDJ rearrangement | CDR3 | P/NP | tissue | # SHM |
|---|---|---|---|---|---|---|---|
| 3315 | Ccnd1/ Ikk2ca | VHB-JH1 | Ighv2-2*02; Ighd2-4*01; Ighj1*01 | GCCAGAACTATGATTACGTACTTCGATGTC | P | F, BMPC | 0 to 1 |
| | | VHB-JH3 | Ighv2-9*02; Ighd2-10*02; Ighj2*01 | GCCGCCCCCTACTATGGTAACGCCTGGTTTGCTTAC | P | F, BMPC | 0 to 1 |
| 3433 | Ccnd1/ Ikk2ca | VHE-JH1 | Ighv5-1*01; Ighd1-2*01; Ighj2*01 | TTGAGACAACATTAGGGGCTAC | NP | SPL, F, BMPC | 1 |
| | | VHF-JH1 | Ighv7-1*01; Ighd1-1*02; Ighj1*03 | GCAAGAGATGATGGTTACTGGTACTTCGATGTC | P | SPL, F, BMPC | 1 to 2 |
| 3443 | Ccnd1/ Ikk2ca | VHE-JH4 | Ighv5-17*02; Ighd2-14*01; Ighj4*01 | GCAACCTACTATAGGTACGACGGCTATGCTTTGGACTAC | P | SPL, F, BMPC | 3 to 4 |
| 3459 | Ccnd1/ Ikk2ca | VHB-JH1 | Ighv2-6-7*01; Ighd2-4*01; Ighj3*01 | GCCAGAGGAGGCAACTATGATTACGACGGGTTTGCTTAC | P | BMPC | 0 |
| | | VHG-JH4 | Ighv9-3*01; Ighd4-1*01; Ighj4*01 | GCAAGACGGGCTGGGACGGAGGCTATGGACTAC | P | BMPC | 0 to 1 |
| | | VHG-JH4 | Ighv9-3*02; Ighd1-1*01; Ighj4*01 | GCAAGAAGGGTTTATTACTACGGTCCTTATGCTATGGACTAC | P | BMPC | 0 to 1 |
| 3439 | Ccnd1/ Ikk2ca | VHF-JH1 | Ighv7-1*02; Ighd4-1*01; Ighj1*01 | GCAAGAGATAACTGGGACTGGTACTTCGATGTC | P | BMPC | 1 |
| 3468 | Ccnd1/ Ikk2ca | VHE-JH4 | Ighv5-6-4*01; Ighd3-3*01; Ighj4*01 | ACAAGAGATCTGGGGACGGAGGGGTATGCTATGGACTAC | P | SPL, BMPC | 1 |
| | | VHB-JH4 | Ighv2-9*02; Ighd2-14*01; Ighj4*01 | GCCAGAGATATGTACGACTATGCTATGGACTAC | P | SPL, BMPC | 0 |
| | | VHE-JH3 | Ighv5-9-1*02; Ighd2-4*01; Ighj3*01 | ACAAGAGATCATGATTACGACGGTTTGCTTAC | NP | SPL, BMPC | 4 |
| | | VHA-JH3 | Ighv1-19*01; Ighd1-1*01; Ighj3*01 | GCCGTATTAATTACTACGGTACTAGAGGGGTTTGCTTAC | P | SPL, BMPC | 15 |

**Table S5**

| Reagents | Company | RR:ID | Catalog-Number | Methods |
|---|---|---|---|---|
| α-B220-BV785 | BioLegend | AB_11218795 | 103245 | Flow Cytometry |
| α-CD19-BV650 | BioLegend | AB_11204087 | 115541 | Flow Cytometry |
| α-CD38-AlexaFluor700 | Invitrogen | AB_657740 | 56-0381-82 | Flow Cytometry |
| α-Fas-PECy7 | BD Biosciences | AB_396768 | 557653 | Flow Cytometry |
| α-IgG1-PE | BD Biosciences | AB_393553 | 550083 | Flow Cytometry |
| α-CD138-APC | BioLegend | AB_10962911 | 142506 | Flow Cytometry |
| α-TACI-PE | BD Biosciences | AB_647234 | 558410 | Flow Cytometry |
| α-Igκ-FITC | BD Biosciences | AB_393527 | 550003 | Immunofluorescence |
| α-Igλ-FITC | BD Biosciences | AB_394854 | 553434 | Immunofluorescence |
| α-CD11c-PE | BioLegend | AB_313776 | 117307 | Immunofluorescence |
| α-CD138-PE | BioLegend | AB_10916119 | 142504 | Immunofluorescence |
| α-CD19-APC | BioLegend | AB_313647 | 115512 | Immunofluorescence |
| α-B220-APC | BioLegend | AB_312997 | 103212 | Immunofluorescence |
| α-mouse IgM-UNLB | Southern Biotech | AB_2794408 | 1070-01 | ELISA |
| α-mouse IgM-BIO | Southern Biotech | AB_2737411 | 1020-08 | ELISA/ELISPOT |
| α-mouse IgG-UNLB | Southern Biotech | AB_2794290 | 1030-01 | ELISA |
| α-mouse IgG-AP | Southern Biotech | AB_2794293 | 1030-04 | ELISA |
| α-mouse IgA-UNLB | Southern Biotech | AB_2314669 | 1040-01 | ELISA |
| α-mouse IgA-BIO | Southern Biotech | AB_2794374 | 1040-08 | ELISA/ELISPOT |
| α-mouse IgG1-BIO | Southern Biotech | AB_2794427 | 1071-08 | ELISPOT |
| α-mouse IgG2a-BIO | Southern Biotech | AB_2794479 | 1080-08 | ELISPOT |
| α-mouse IgG2b-BIO | Southern Biotech | AB_2794523 | 1090-08 | ELISPOT |
| α-mouse IgG2c-BIO | Southern Biotech | AB_2794463 | 1078-08 | ELISPOT |
| α-mouseIgG3-BIO | Southern Biotech | AB_2794575 | 1100-08 | ELISPOT |
| α-mouse Igκ-UNLB | Southern Biotech | AB_2737431 | 1050-01 | ELISPOT |
| α-mouse Igλ-UNLB | Southern Biotech | AB_2794389 | 1060-01 | ELISPOT |
| Streptavidin-AP | MERCK | | 11089161001 | ELISA |
| BCIP/NBT | Promega | | S381C | ELISPOT |
| α-mouse IgM | SIGMA | AB_260700 | M 8644 | SPEP-Immunofixation |
| α-mouse IgG | SIGMA | AB_260466 | M 1397 | SPEP-Immunofixation |

| | | | | |
|---|---|---|---|---|
| α-mouse IgA | SIGMA | AB_260464 | M 1272 | SPEP-Immunofixation |
| α-mouse Igκ | NOVUS Biologicals | AB_525186 | NB7546 | SPEP-Immunofixation |
| α-mouse Igλ | NOVUS Biologicals | AB_525259 | NB7550 | SPEP-Immunofixation |
| α-WHSC1/NSD2 | abcam | AB_1310816 | ab75359 | Immunoblot |
| α-Cyclin D1 | Cell Signaling | AB_2228523 | 2922 | Immunoblot |

**Table S6**

| module | origin | # genes with mouse ensemble annotation |
|---|---|---|
| MM_signature_UP | Zhan et al., 2002; Table 5 + Lopez-Corral et al., 2014; Table S4 | 49 (38 + 11) |
| MM_signature_DOWN | Zhan et al., 2002; Table 4 + Lopez-Corral et al., 2014; Table S4 | 81 (26 + 55) |
| Gutierrez_WM | Gutierrez et al., 2007; Table 4 | 33 |
| Gutierrez_MM | Gutierrez et al., 2007; Table 3 | 8 |
| Bergsagel_11q13_UP | Bergsagel et al., 2005; Suppl. Table "MM 11q13 genes" | 342 |
| Bergsagel_MMSET_UP | Bergsagel et al., 2005; Suppl. Table "MM 4p16 genes" | 230 |
| Bergsagel_maf_UP | Bergsagel et al., 2005; Suppl. Table "MM maf genes" | 236 |
| Bergsagel_D1_UP | Bergsagel et al., 2005; Suppl. Table "MM D1 genes" | 410 |
| Bergsagel_D2_UP | Bergsagel et al., 2005; Suppl. Table "MM D2 genes" | 150 |
| Zhan_CD-1_UP | Zhan et al., 2006; Table S2; subgroup CD-1 | 34 |
| Zhan_CD-1_DOWN | Zhan et al., 2006; Table S3; subgroup CD-1 | 31 |
| Zhan_CD-2_UP | Zhan et al., 2006; Table S2; subgroup CD-2 | 35 |
| Zhan_CD-2_DOWN | Zhan et al., 2006; Table S3; subgroup CD-2 | 33 |
| Zhan_MMSET_UP | Zhan et al., 2006; Table S2; subgroup MS | 33 |
| Zhan_MMSET_DOWN | Zhan et al., 2006; Table S3; subgroup MS | 28 |
| Zhan_MF_UP | Zhan et al., 2006; Table S2; subgroup MF | 35 |
| Zhan_MF_DOWN | Zhan et al., 2006; Table S3; subgroup MF | 31 |
| Zhan_HP_UP | Zhan et al., 2006; Table S2; subgroup HP | 31 |
| Zhan_HP_DOWN | Zhan et al., 2006; Table S3; subgroup HP | 34 |
| Zhan_LB_UP | Zhan et al., 2006; Table S2; subgroup LB | 35 |
| Zhan_LB_DOWN | Zhan et al., 2006; Table S3; subgroup LB | 25 |
| Zhan_PR_UP | Zhan et al., 2006; Table S2; subgroup PR | 34 |
| Zhan_PR_DOWN | Zhan et al., 2006; Table S3; subgroup PR | 59 |
| Broyl_CD-1_UP | Broyl et al., 2010; Table S5; Cluster CD1 top up-regulated (50) | 29 |
| Broyl_CD-1_DOWN | Broyl et al., 2010; Table S5; Cluster CD1 top down-regulated (9) | 5 |

| | | |
|---|---|---|
| Broyl_CD-2_UP | Broyl et al., 2010; Table S5; Cluster CD2 top up-regulated (50) | 24 |
| Broyl_CD-2_DOWN | Broyl et al., 2010; Table S5; Cluster CD2 top down-regulated (50) | 20 |
| Broyl_CTA_UP | Broyl et al., 2010; Table S5; Cluster CTA top up-regulated (50) | 30 |
| Broyl_CTA_DOWN | Broyl et al., 2010; Table S5; Cluster CTA top down-regulated (50) | 31 |
| Broyl_NFkB_UP | Broyl et al., 2010; Table S5; Cluster NFkB top up-regulated (50) | 28 |
| Broyl_NFkB_DOWN | Broyl et al., 2010; Table S5; Cluster NFkB top down-regulated (50) | 39 |
| Broyl_HY_UP | Broyl et al., 2010; Table S5; Cluster HY top up-regulated (50) | 28 |
| Broyl_HY_DOWN | Broyl et al., 2010; Table S5; Cluster HY top down-regulated (50) | 28 |
| Broyl_PRL3_UP | Broyl et al., 2010; Table S5; Cluster PRL3 top up-regulated (18) | 11 |
| Broyl_PRL3_DOWN | Broyl et al., 2010; Table S5; Cluster PRL3 top down-regulated (9) | 6 |
| Broyl_PR_UP | Broyl et al., 2010; Table S5; Cluster PR top up-regulated (50) | 34 |
| Broyl_PR_DOWN | Broyl et al., 2010; Table S5; Cluster PR top down-regulated (50) | 31 |
| Broyl_MF_UP | Broyl et al., 2010; Table S5; Cluster MF top up-regulated (50) | 29 |
| Broyl_MF_DOWN | Broyl et al., 2010; Table S5; Cluster MF top down-regulated (50) | 24 |
| Broyl_MMSET_UP | Broyl et al., 2010; Table S5; Cluster MS top up-regulated (50) | 25 |
| Broyl_MMSET_DOWN | Broyl et al., 2010; Table S5; Cluster MS top down-regulated (50) | 26 |
| Broyl_Myeloid_UP | Broyl et al., 2010; Table S5; Cluster Myeloid top up-regulated (50) | 17 |
| Broyl_Myeloid_DOWN | Broyl et al., 2010; Table S5; Cluster Myeloid top down-regulated (40) | 28 |

**References (35-47)**

35. T. Sommermann, T. Yasuda, J. Ronen, T. Wirtz, T. Weber, U. Sack, R. Caeser, J. Zhang, X. Li, V. Trung Chu, A. Jauch, K. Unger, D. J. Hodson, A. Akalin, K. Rajewsky, A. con-, Functional interplay of Epstein-Barr virus oncoproteins in a mouse model of B cell lymphomagenesis. *PNAS*. **117**, 14421–14432 (2020).
36. Z. Hao, K. Rajewsky, Homeostasis of Peripheral B Cells in the Absence of B Cell Influx from the Bone Marrow. *J. Exp. Med*. **194**, 1151–1163 (2001).
37. J. P. DiSanto, W. Müller, D. Guy-Grand, A. Fischer, K. Rajewsky, Lymphoid development in mice with a targeted deletion of the interleukin 2 receptor g chain. *PNAS*. **92**, 377–381 (1995).
38. K. Schmidt, U. Sack, R. Graf, W. Winkler, O. Popp, P. Mertins, T. Sommermann, C. Kocks, K. Rajewsky, B-Cell-Specific Myd88 L252P Expression Causes a Premalignant Gammopathy Resembling IgM MGUS. *Front. Immunol.* **11** (2020), doi:10.3389/fimmu.2020.602868.
39. M. Peitz, K. Pfannkuche, K. Rajewsky, F. Edenhofer, Ability of the hydrophobic FGF and basic TAT peptides to promote cellular uptake of recombinant Cre recombinase: A tool for efficient genetic engineering of mammalian genomes. *PNAS*. **99**, 4489–4494 (2002).
40. T. Nojima, K. Haniuda, T. Moutai, M. Matsudaira, S. Mizokawa, I. Shiratori, T. Azuma, D. Kitamura, In-vitro derived germinal centre B cells differentially generate memory B or plasma cells in vivo. *Nat. Commun.* **2**, 1–11 (2011).
41. D. Krappmann, F. Emmerich, U. Kordes, E. Scharschmidt, B. Doè Rken, C. Scheidereit, Molecular mechanisms of constitutive NF-kB/Rel activation in Hodgkin/ Reed-Sternberg cells. *Oncogene*. **18**, 943–953 (1999).
42. S. L. Rosales, S. Liang, I. Engel, B. J. Schmiedel, M. Kronenberg, P. Vijayanand, G. Seumois, R. L. Reinhardt, Ed. (Springer New York, New York, NY, 2018; https://doi.org/10.1007/978-1-4939-7896-0_21), pp. 275–302.
43. R. Patro, G. Duggal, M. I. Love, R. A. Irizarry, C. Kingsford, Salmon provides fast and bias-aware quantification of transcript expression. *Nat. Methods*. **14**, 417–419 (2017).
44. C. Soneson, M. I. Love, M. D. Robinson, Differential analyses for RNA-seq: transcript-level estimates improve gene-level inferences. *F1000Research*. **4**,

1–19 (2016).
45. M. I. Love, W. Huber, S. Anders, Moderated estimation of fold change and dispersion for RNA-seq data with DESeq2. *Genome Biol.* **15**, 1–21 (2014).
46. J. Weiner, Feature Set Enrichment Analysis for Metabolomics and Transcriptomics (2020), (available at https://cran.r-project.org/package=tmod).
47. G. Yu, enrichplot: Visualization of Functional Enrichment Result. (2021), (available at https://yulab-smu.top/biomedical-knowledge-mining-book/).
